# Noncanonical mRNA decay by the endoplasmic-reticulum stress sensor IRE1α promotes cancer-cell survival

**DOI:** 10.1101/2021.03.16.435520

**Authors:** Adrien Le Thomas, Elena Ferri, Scot Marsters, Jonathan M. Harnoss, Zora Modrusan, Weihan Li, Joachim Rudolph, Weiru Wang, Thomas D. Wu, Peter Walter, Avi Ashkenazi

## Abstract

Eukaryotic IRE1 mitigates endoplasmic-reticulum (ER) stress by orchestrating the unfolded-protein response (UPR). IRE1 spans the ER membrane, and signals through a cytosolic kinase-endoribonuclease module. The endoribonuclease generates the transcription factor XBP1s by intron excision between similar RNA stem-loop endomotifs, and depletes select cellular mRNAs through regulated IRE1-dependent decay (RIDD). Paradoxically, mammalian RIDD seemingly targets only mRNAs with XBP1-like endomotifs, while in flies RIDD exhibits little sequence restriction. By comparing nascent and total IRE1α-controlled mRNAs in human breast cancer cells, we discovered not only canonical endomotif-containing RIDD substrates, but also many targets lacking recognizable motifs—degraded by a process we coin RIDDLE, for RIDD lacking endomotif. IRE1α displayed two basic endoribonuclease modalities: endomotif-specific cleavage, minimally requiring dimers; and endomotif-independent promiscuous processing, requiring phospho-oligomers. An oligomer-deficient mutant that did not support RIDDLE failed to rescue cancer-cell viability. These results link IRE1α oligomers, RIDDLE, and cell survival, advancing mechanistic understanding of the UPR.

## INTRODUCTION

The ER mediates folding of newly synthesized secretory and membrane proteins. Excess folding demand leads to ER accumulation of misfolded proteins, causing ER stress. This engages an intracellular signaling network, dubbed the unfolded protein response (UPR), to reestablish homeostasis^1–3^. The mammalian UPR entails three ER-transmembrane proteins: IRE1α, PERK, and ATF6, which coordinate adaptive changes to expand ER capacity while abating ER load^1,3,4^. If adaptation fails, the UPR triggers apoptotic cell death^4–6^. UPR dysregulation contributes to several diseases^7–11^. Cancer cells often leverage the UPR, including IRE1α, to circumvent ER stress and maintain malignant growth^10,12–17^. Better mechanistic understanding would help elucidate the UPR’s role in disease, and advance its potential for medical translation.

IRE1α comprises ER-lumenal and transmembrane domains, and a cytosolic kinase-endoribonuclease (KR) module^18,19^. Unfolded-protein sensing by the lumenal domain drives IRE1α homodimerization, kinase-mediated *trans*-autophosphorylation, and endoribonuclease activation^19–24^. The RNase produces the transcription factor spliced X-box binding protein 1 (XBP1s), and depletes multiple cellular mRNAs through regulated IRE1-dependent decay (RIDD)^25–27^. XBP1s-target genes support protein folding and ER-associated degradation (ERAD)^28^. IRE1α cleaves unspliced XBP1<u>u</u>mRNA at two stem-loop endomotifs, removing a 26-nt intron^29–31^, and RtcB ligates the severed exons, generating XBP1s^32–34^. XBP1u cleavage at each splice site requires an energetically stable stem, as well as a 7-nt consensus sequence CNG|CAGC within the loop, with scission between G and C in the third and fourth positions^34,35^.

RIDD remains puzzling^27,36^. In the budding yeast *S. cerevisiae*, IRE1 triggers non-conventional mRNA splicing of the *XBP1* ortholog *HAC1*, yet lacks RIDD activity^26^. Conversely, in the fission yeast *S. pombe*, IRE1 performs RIDD, which targets a UG|CU core motif within variably sized stem-loop structures, but *HAC1* mRNA is absent^37^. In the fruit fly *D. melanogaster*, RIDD primarily targets ER-bound mRNAs, with minimal sequence and unknown structure restriction^26,38,39^. By contrast, in mammals, known RIDD substrates seem to possess and require for cleavage an XBP1-like consensus loop sequence CNGCAGN, enclosed by a stable stem^40–43^. Besides ER load, mammalian RIDD regulates additional cellular functions, including triglyceride and cholesterol metabolism^44^; apoptosis signaling through DR5^45–47^; protective autophagy via BLOC1S1 (BLOS1)^48^; and DNA repair through Ruvbl1^49,50^. A canonical stem-loop endomotif is necessary but not sufficient to predict mammalian RIDD, while translational stalling can enhance mRNA depletion^41^. Although other mechanisms, such as NO-GO decay and the cytosolic exosome, have been implicated in completing degradation after endomotif-directed mRNA cleavage by IRE1^51^, it is unknown whether IRE1 itself can conduct full RNA digestion. Autophosphorylation supporting IRE1 oligomerization^22,45,50,52^ and the resulting higher-order assemblies may affect RNase output^21,22,53–57^. However, the specific requirements for endomotif-restricted *vs*. non-restricted IRE1 RNase activity remain ill-defined, and the significance of the latter in cells of higher metazoans remains elusive.

To investigate the scope of RIDD in a human cancer cell line, we took a dual whole-genome RNA sequencing approach that distinguishes total cellular mRNAs from nascent transcripts. Surprisingly, in ER-stressed human MDA-MB-231 breast cancer cells, IRE1α depleted not only canonical endomotif-containing RIDD substrates, but also multiple targets of “RIDD lacking <u>e</u>ndomotif” (RIDDLE). By isolating homotypic complexes of the human IRE1α kinase-endoribonuclease module in monomeric, dimeric, or oligomeric form, we discovered that RIDD and RIDDLE reside in two distinct RNase modalities: endomotif-directed cleavage, minimally requiring IRE1α dimers; and endomotif-independent cleavage, requiring phospho-oligomers. Indeed, an IRE1α mutant that could form dimers but not oligomers selectively lost the RIDDLE modality. This mutant failed to rescue cancer-cell viability upon depletion of wildtype IRE1α, demonstrating biological significance of IRE1α oligomers and RIDDLE.

## RESULTS

### Integrated RNAseq and GROseq identify mRNA targets of IRE1α-dependent decay

We reasoned that subtracting nascent-transcript alterations from global changes in mRNA abundance would help distinguish mRNA decay from diminished transcription. Accordingly, to examine mRNAs subject to IRE1-dependent decay, we applied two parallel RNA sequencing approaches: (1) classical RNAseq, which interrogates the steady-state transcriptome; and (2) global nuclear run-on sequencing (GROseq)^58^, which interrogates the nascent transcriptome. We determined dependency on IRE1α and ER stress by treating human MDA-MB-231 breast cancer cells, harboring homozygous wildtype (WT) or CRISPR/Cas9 knockout (KO) *IRE1*α alleles^14^, with the classical ER stressor thapsigargin (Tg).

Most global-transcript levels did not change under these conditions (Fig. S1a). We identified 54 mRNAs as potential RIDD substrates: these displayed an IRE1α-dependent, ER stress-induced decrease in abundance of at least 1.4-fold as measured by RNAseq, without corresponding declines in transcription as measured by GROseq (Table S1). We illustrate regulation of 8 of these mRNAs, including the known RIDD targets CD59 and DGAT2, as well as several novel ones, namely, TGOLN2, GBA, SNN, SIX2, TNFAIP18L1, and MFAP2 (Fig. 1a). By contrast, other mRNAs showed more complex behaviors: SCARA3 and PRICKLE2 mRNA levels declined both as measured by RNAseq and GROseq, exemplifying a class of mRNAs that decrease in abundance but may not be directly cleaved by IRE1, consistent with other results^25,41^. CD59 and GBA showed down-regulation via both RIDD and diminished transcription; and MFAP2 was upregulated by ER stress while being suppressed via IRE1α. Individual mRNAs displayed different decay rates, with some—such as TNFAIP8L1, SNN, and SIX2—showing a significant lag in depletion after Tg addition (Fig. 1a-b and Fig. S1b). Bioinformatic analysis suggested a notable frequency of hits in annotated categories of cell morphology, and cell death or survival (Table S2). Many lacked a signal peptide or anchor (Table S1), consistent with modulation of diverse cellular functions^45–47^.

**Figure 1.**
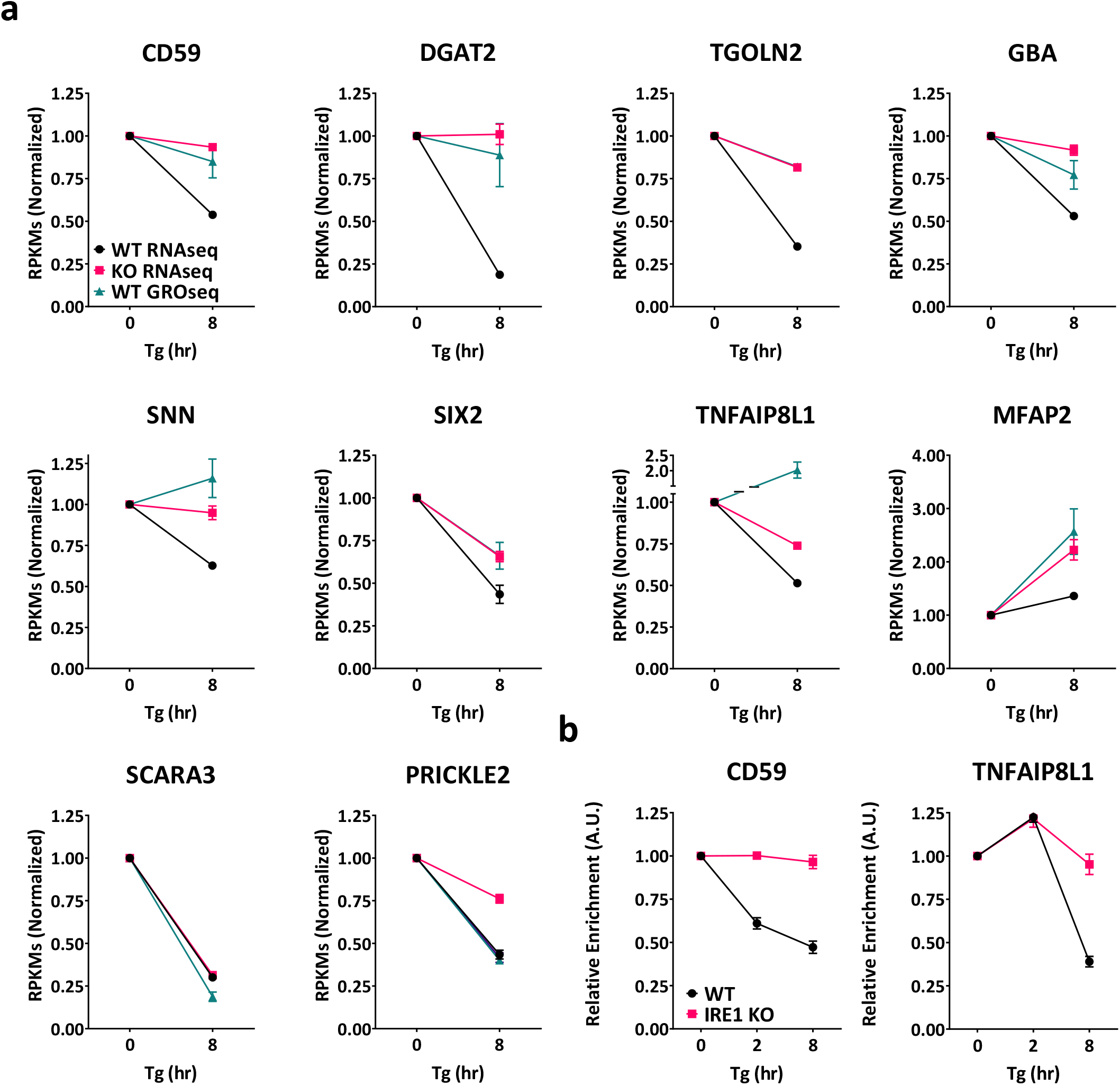
Integrative RNAseq and GROseq analyses identify human RIDD and RIDDLE targets. (**a**) Mean RPKM values for various examples of IRE1α RNase targets from the RNAseq and GROseq datasets in WT and IRE1α KO MDA-MB-231 cells before and after ER-stress induction by Tg (100 nM). (**b**) Kinetic RT-qPCRs analysis of CD59 and TNFAIP8L1 transcripts in WT and IRE1 KO MDA-MB-231 cells, before and after ER-stress induction by Tg (100 nM) for 2 and 8 hours.

As expected, GROseq detected over 300 mRNAs that displayed IRE1α-dependent transcriptional upregulation by ER stress, enriched in gene-ontology of ER stress, ER-to-Golgi vesicle transport, IRE1-mediated UPR, N-linked glycosylation, and ERAD (Table S3). Several mRNAs represented known XBP1s transcriptional targets (Fig. S1c). GROseq also covered the genomic region encoding the XBP1u intron (Fig. S1d), providing further methodological validation.

### IRE1α displays two distinct endoribonuclease modalities

To explore molecular features that might govern RNase activity, we purified recombinant human IRE1α kinase-endoribonuclease (IRE1-KR) proteins in unphosphorylated (0P) or fully phosphorylated (3P) states (Fig. S2a). IRE1-KR-0P efficiently cleaved an XBP1u-based T7 RNA polymerase transcript, at both of the known stem-loop endomotifs: processing produced ∼500-nt and ∼350-nt fragments, corresponding to the 5’ and 3’ exons; and the 26-nt intron (Fig. 2a, Fig. S2b,c). Scrambling either loop sequence prevented cleavage, while inserting 43-nt or 50-nt random sequences between the splice sites proportionally shifted the resulting bands (Fig. S2b,c). Thus, IRE1-KR-0P performs precise cleavage of XBP1u RNA at the two consensus sites, in keeping with other data^22,29,31^.

**Figure 2.**
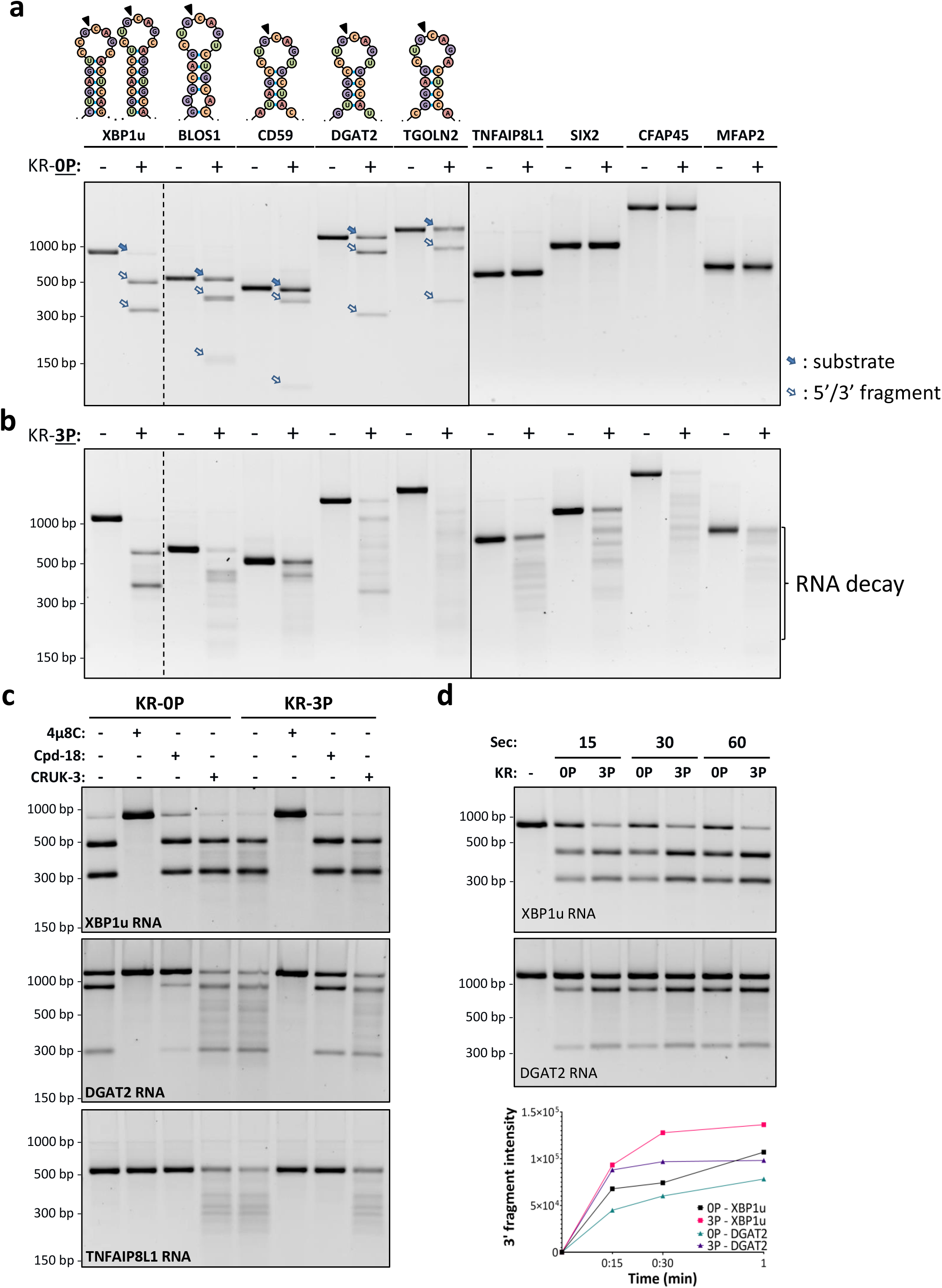
Phosphorylation state of IRE1α affects RNase modality. In vitro-generated T7-RNAs were incubated with purified recombinant IRE1α protein (residues G547-L977) comprising the kinase-endoribonuclease module (IRE1-KR), in non-phosphorylated (KR-0P) (**a**) or fully phosphorylated (KR-3P) (**b**) form, and RNA products were analyzed by agarose gel electrophoresis. Where shown, solid arrows indicate RNA substrates; open arrows mark RNA cleavage products, representing the 5’ and 3’ fragments around the cleavage site. Specific RNA endomotifs are depicted on top for respective targets. (**c**) KR-0P and KR-3P digestions of XBP1, DGAT2, and TNFAIP8L1 transcripts in the presence of the IRE1α RNase inhibitor 4μ8C, the IRE1α kinase-based RNase inhibitor Compound 18 (Cpd-18), or the IRE1α kinase-based RNase activator Compound 3 (CRUK-3). (**d**) KR-3P digestion of XBP1u and DGAT2 mRNA at shorter duration. Intensities of the 3’ RNA fragment were quantified for XBP1u and DGAT2 using GelQuantNET from the RNA digestion agarose gels shown above.

Next, we examined the capacity of IRE1-KR-0P to process 8 of the 54 potential RIDD targets we identified, including 3 established and 5 newly uncovered ones (Fig. 2a). IRE1-KR-0P performed single-site cleavage of 4 of the RNAs, encoding BLOS1, CD59, DGAT2, and TGOLN2. Each of these contains an XBP1-like stem-loop endomotif, having the core consensus sequence CNGCAGN within a projected stem-loop secondary structure; the 5’ and 3’ fragments produced by IRE1-KR-0P for each RNA agreed in size with the location of the stem-loop endomotif (Fig. 2a and Table S4). Compared to XBP1u RNA, processing of the latter transcripts left more RNA substrate intact at the timepoint analyzed, indicating generally slower or less efficient reactions. Scrambling the loop sequence of CD59 and DGAT2 prevented cleavage by IRE1-KR-0P (Fig. S2d), confirming its endomotif-restricted endoribonuclease activity.

Surprisingly, under the same reaction conditions, IRE1-KR-0P failed to cleave the other 4 RNAs, encoding TNFAIP18L1, SIX2, CFAP45, and MFAP2 (Fig. 2a). Resistance of these RNAs to cleavage correlated with their lack of a robust canonical stem-loop endomotif, as described below.

IRE1-KR-3P produced the same XBP1u RNA cleavage fragments as did IRE1-KR-0P; however, IRE1-KR-3P generated additional XBP1u fragments, visible as a faint smear (Fig. 2b), suggesting that it can catalyze further RNA decay. Uniquely, IRE1-KR-3P also cleaved into multiple fragments each of the 8 RNA substrates, including those that resisted cleavage by IRE1-KR-0P. Since these more promiscuous cleavages occur outside the endomotif, we categorize them as RIDDLE activity.

To ascertain dependence on IRE1α, we included the IRE1α RNase-directed inhibitor 4μ8c^59^, which completely prevented RNA cleavage by both IRE1-KR-0P and IRE1-KR-3P (Fig. 2c). In contrast, the IRE1α kinase-directed RNase inhibitor Compound 18^15,60^ blocked RNA decay, but not endomotif cleavage; and the kinase-based IRE1α RNase activator CRUK-3^61^ endowed IRE1-KR-0P with an IRE1-KR-3P-like ability to cleave RNA in both modalities. Upon processing by IRE1-KR-3P, endomotif-mutated XBP1u and CD59 RNA substrates were more stable than WT counterparts (Fig. S2d), suggesting that endomotif-based cleavage can prime mRNA for RIDDLE, which then leads to further decay of the initial fragments. Kinetic analyses suggested faster endomotif cleavage by IRE1-KR-3P than IRE1-KR-0P, evident by swifter generation of 3’ products from substrate RNAs (Fig. 2d and Fig S2e), and more rapid scission of a short synthetic hairpin endomotif (Fig. S2f). These results demonstrate that IRE1α can switch between two different endoribonuclease modalities: (1) endomotif-restricted activity, which mediates both XBP1u intron excision and canonical RIDD; (2) endomotif-independent activity, which mediates RIDDLE, as well as further degradation after canonical endomotif recognition. IRE1-KR-0P supports the first modality but not the second, whereas IRE1-KR-3P conducts both.

Combining the data for the above 8 RNAs with empirical results for 4 additional targets, *i*.*e*., PIGQ, BMP4, BCAM, and SNN (Fig. S2g), together with 29 earlier characterized human or mouse RIDD substrates^27,43^, we developed a new computational algorithm, dubbed gRIDD, which accounts for all of these verified RIDD targets, and determines for any given mRNA the presence of any canonical stem-loop endomotifs (see Supplemental Methods for a detailed description). This algorithm takes into account features that include: (1) conformity of the loop sequence to the consensus; (2) loop length and stem stability in context of the 55-60 flanking bases; (3) number of paired bases and any unpaired bases in the stem. To validate gRIDD, we analyzed a test-set of 4 additional transcripts from our screen. Two were found computationally to possess a canonical stem-loop endomotif, with either an exact match to (GBA) or a single nt variation from (WT1) the consensus loop sequence; accordingly, both IRE1-KR-0P and IRE1-KR-3P should cleave these latter RNAs. Another test RNA (CCDC69) has weak endomotif that fails gRIDD criteria, while a fourth one (AIM2) lacks an endomotif altogether; accordingly only IRE1-KR-3P should cleave these latter RNAs. Supporting gRIDD’s accuracy, all test RNAs displayed the predicted cleavage characteristics (Fig. S2h). Providing further validation, the known RIDD targets murine ANGPTL3^44^, murine SUMO2, human SUMO3^42^, and human DR5^47^ met the algorithm’s criteria, whereas 10 mRNAs previously excluded from RIDD^41^ did not.

Of the 54 mRNAs we identified here (Table S4), gRIDD mapped 30 transcripts as possessing a canonical stem-loop endomotif (RIDD), including 22 with an exact and 8 with an acceptably variant, consensus loop sequence. In addition, gRIDD mapped 24 transcripts as lacking an endomotif (RIDDLE), including 15 with one or more sub-par stem-loop endomotif that failed the algorithm’s criteria, and another 9 that had no discernable endomotif. We empirically confirmed RNAs from each of these subclasses.

### RIDDLE is promiscuous in substrate recognition yet non-random

To test the uniformity of RNA decay by IRE1-KR-3P, we performed 3 independent RNA digestions of DGAT2 and TNFAIP8L1. Strikingly, IRE1-KR-3P generated the same banding pattern in all 3 cases, demonstrating that—although the processing appeared more promiscuous than endomotif-directed cleavage—it entailed consistent, non-random fragmentation (Fig. 3a). To search for underlying sequence requirements, we subjected TNFAIP8L1 RNA to cleavage by IRE1-KR-3P, resolved the products by agarose gel electrophoresis, and extracted them for Sanger sequencing. Overall, 60 of the 69 reads thus obtained indicated cleavage at GC sites, with infrequent cleavage at other sites (Fig. 3b). Alignment along the TNFAIP8L1 RNA indicated two relatively enriched cleavage locations, S1 (21/69) and S2 (11/69), both at GC sites (Fig. 3c). The sequences surrounding these two GC sites did not meet gRIDD’s stem-loop endomotif criteria. Regardless, replacing GC by TA at S1 or S2 abolished or diminished the corresponding cleavage product (Fig. 3d), confirming site selectivity. Likewise, analysis of DGAT2 RNA also revealed a preponderance of GC cleavages (34/61 reads, excluding the endomotif), and mutation of the most prevalent site precluded the corresponding product (Fig. S3 a-c). Thus, although RIDDLE is less restricted, it appears to favor GC sites.

**Figure 3.**
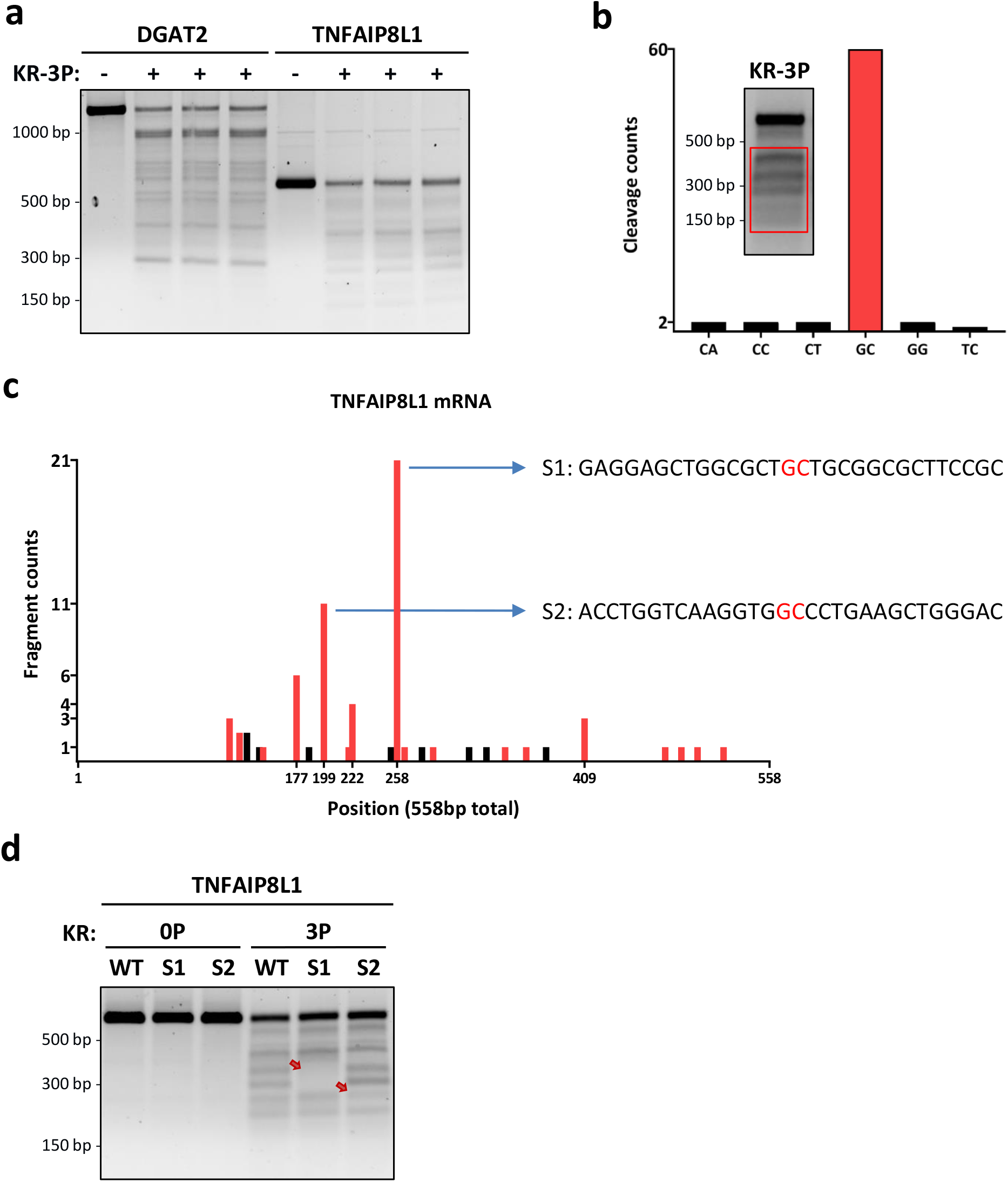
RIDDLE is relatively unrestricted by substrate sequence but non-random. (**a**) Comparative KR-3P digestion of DGAT2 and TNFAIP8L1 transcripts performed in three independent experiments. (**b**) Amount of RNA fragments sequenced whose 3’ end leads to the cleaved nt pair designated on the x axis. The first nt in the pair represents the last sequenced nt from the RNA fragment, while the second shows the subsequent base in the RNA sequence. Inset: red box indicates the portion of the gel that was extracted for Sanger sequencing. (**c**) Mapping of the last base pair (3’ end) from each individual RNA fragment sequenced within the TNFAIP8L1 mRNA. Red bars indicate cleavage sites between a GC nt pair. (**d**) RNA digestions of WT TNFAIP8L1 and TNFAIP8L1 mutated at locations S1 and S2. The red arrows indicate change in banding pattern as compared to WT.

### Phospho-oligomeric state governs IRE1α’s endoribonuclease modality

While activation of the IRE1α RNase requires homodimerization, the importance of higher-order assembly is unclear^56,62^. To directly investigate the role of such assembly, we covalently tethered IRE1-KR protomers by chemical crosslinking and studied their RNase modalities. Immunoblot analysis revealed that IRE1-KR-0P was primarily monomeric yet formed some detectable dimers in a concentration-dependent manner; in contrast, IRE1-KR-3P formed not only more prominent concentration-dependent dimers, but also oligomers consistent with relative masses of potential tetramers and hexamers (Fig. 4a). These results agree with other evidence that phosphorylation of IRE1α promotes dimerization and stabilizes higher-order oligomerization^22,52,63,64^.

**Figure 4.**
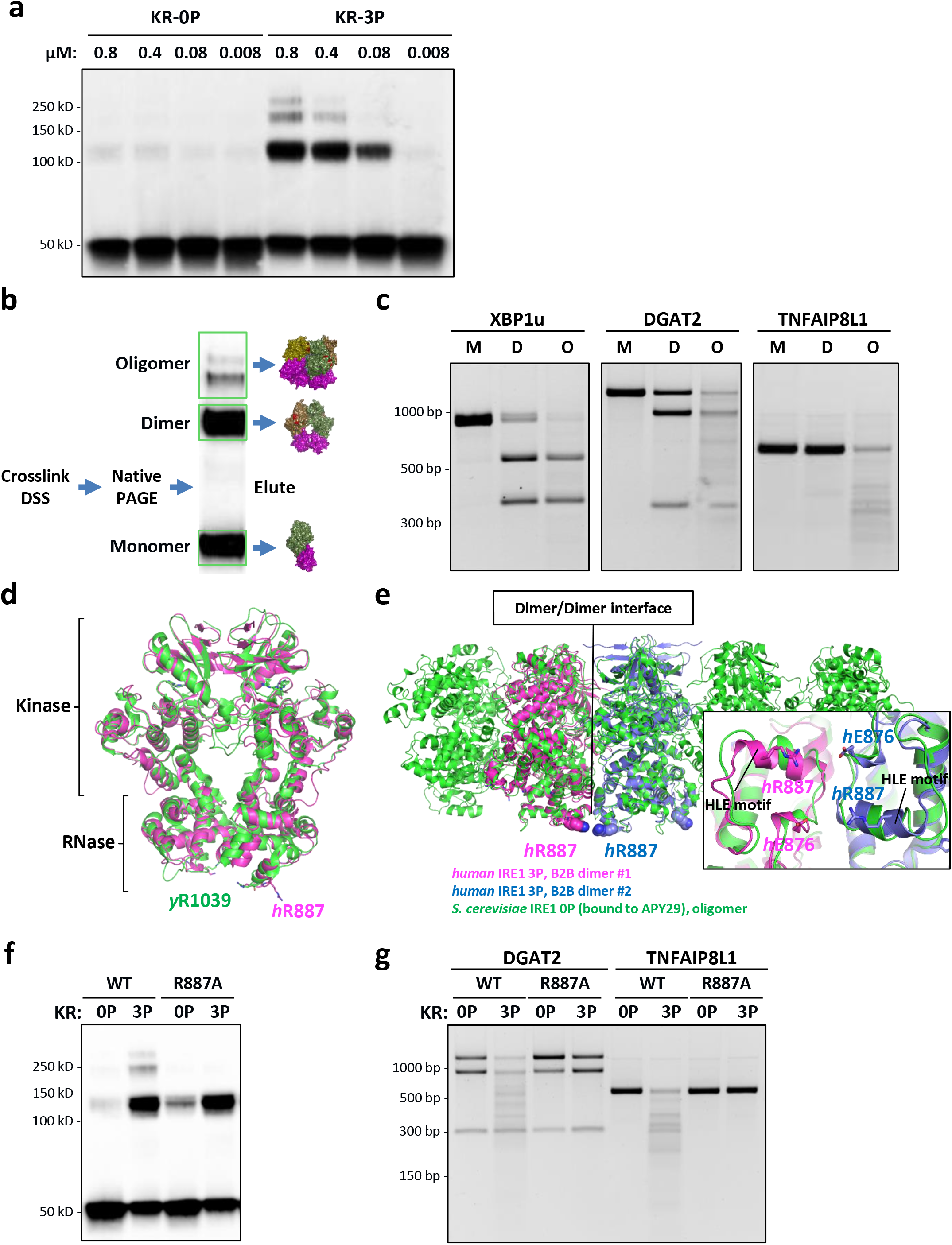
Phospho-oligomeric state governs IRE1α’s RNase modality. (**a**) DSS crosslinking of IRE1-KR-0P andIRE1-KR-3P at various concentrations. (All lanes have the same final amount of protein loaded.) (**b**) Cartoon depicting the procedure to extract KR-3P fractions before incubation with T7 RNAs. (**c**) Digestion of XBP1u, DGAT2, and TNFAIP8L1 RNA by isolated fractions of IRE1-KR-3P: M, monomer; D, dimer; and O, oligomer. (**d**) WT structure of IRE1α dimer with the residues of interest highlighted: alignment of yeast B2B IRE1α (PDB ID: 3FBV, green) and human B2B IRE1α (PDB ID: 6W3C, magenta). (**e**) Alignment of yeast oligomeric IRE1α (PDB ID: 3FBV) and two dimers of human IRE1α (PDB ID: 6W3C). Inset: R887-E876 spatial arrangement based on the alignment. (**f**) DSS crosslinking of purified recombinant IRE1-KR-3P WT and R887A protein. (**g**) Digestion of DGAT2 and TNFAIP8L1 RNA by IRE1-KR-3P WT and R887A.

IRE1-KR-3P retained RNase activity after crosslinking, evident by cleavage of XBP1u, DGAT2, and TNFAIP8L1 RNAs (Fig. S4a). To examine RNase modality, we resolved crosslinked IRE1-KR-3P complexes into monomers, dimers, and oligomers by native polyacrylamide gel electrophoresis (Fig. 4b), excised and eluted each complex, and reacted it with RNA substrate (Fig. 4c). Whereas IRE1-KR-3P monomers were inactive as expected, dimers excised the XBP1u intron at both splice sites, and cleaved the DGAT2 RNA endomotif; however, they failed to further degrade DGAT2 RNA appreciably, nor did they cleave TNFAIP8L1 RNA. In contrast, IRE1-KR-3P oligomers cleaved XBP1u more completely, while substantially degrading both DGAT2 and TNFAIP8L1 RNAs. Upon prolonged incubation, IRE1-KR-3P oligomers achieved complete RNA degradation, as exemplified for DGAT2 RNA (Fig. S4b). Thus, whereas endomotif-directed cleavage minimally requires dimers, RIDDLE requires higher-order oligomerization. Supporting this conclusion, Compound 18, which selectively blocked endomotif-independent cleavage by IRE1-KR-3P (Fig. 2d), inhibited IRE1-KR-3P oligomerization but not dimerization (Fig. S4c). Moreover, CRUK-3, which enhanced endomotif-directed cleavage and endomotif-independent degradation (RIDDLE) by IRE1-KR-0P (Fig. 2d), congruently augmented IRE1-KR-0P oligomerization (Fig. S4c). Mutational analysis confirmed that at least two of the three phosphorylation sites within the kinase activation loop of IRE1α (serine 724, 726, 729) were required for enhanced dimer formation, higher-order oligomerization, and the corresponding RNase modality. Of note, the triply phosphorylated protein showed a markedly better capacity to oligomerize and to perform RNA decay (Fig. S4d).

To seek further mechanistic insight, we screened for human IRE1α mutations that might disrupt oligomerization, by testing several amino acid positions previously studied with yeast IRE1^62,64^. One variant—R887A—proved particularly useful. Arginine 887 resides in the RNase domain within a helix loop element (HLE), shown to be important for binding and cleavage of *HAC1* mRNA in *S. cerevisiae*^64^. As such, R887 does not stabilize IRE1’s so-called back-to-back (B2B) dimer interface (Fig. 4d). However, structural overlay of human B2B dimers (PDB ID: 6W3C) onto oligomeric *S. cerevisiae* IRE1 (PDB ID: 3FBV) places R887 at the interface of two B2B dimers within the projected human oligomer (Fig. 4e). Importantly, although IRE1-KR-R887A dimerized, it failed to form higher-order oligomers, regardless of phosphorylation (Fig. 4f). Compared to WT IRE1-KR-0P, unphosphorylated R887A performed endomotif-directed cleavage of DGAT2 and did not appreciably degrade DGAT2 or TNFAIP8L1 RNAs (Fig. 4g). Strikingly, phosphorylated IRE1-KR-R887A retained endomotif-directed DGAT2 cleavage, but unlike WT IRE1-KR-3P, it failed to degrade DGAT2 and TNFAIP8L1 RNAs. These loss-of-function results strongly validate the conclusion that RIDDLE depends on higher-order phospho-oligomers of IRE1α.

### Oligomer-deficient mutant IRE1α fails to rescue cancer-cell viability

To extend the principles gleaned from our studies of IRE1-KR *in vitro* to full-length IRE1α in cells, we devised a functional complementation strategy: We stably transfected shRNA-resistant cDNA expression plasmids encoding GFP-tagged^56^ WT or IRE1α-R887A into MDA-MB-231 cells harboring doxycycline (Dox)-inducible shRNAs against IRE1α, and isolated GFP-positive transfectants by cell sorting. As expected, each ectopic protein expressed independent of Dox-inducible depletion of endogenous IRE1α (Fig. 5a). The transgenic proteins migrated at a higher molecular mass, consistent with their GFP tagging, and showed elevated expression relative to endogenous IRE1α. RT-qPCR analysis after a 72-hr Dox-induced depletion of intrinsic IRE1α demonstrated that the ectopic WT and mutant variants supported a comparable fold-induction of XBP1s upon ER stress (Fig 5b), suggesting similar capacity for endomotif-directed cleavage. To monitor RNase modality toward RIDD targets, we designed two specific RT-qPCR primer pairs for CD59 or TGOLN2: One encompasses the endomotif and therefore measures endomotif-directed cleavage (RIDD); the other covers the mRNA’s 3’ end and hence detects decay (RIDDLE; schematized in Fig. S5a). Although both WT and R887A mediated endomotif-directed cleavage of both CD59 and TGOLN2 mRNAs, only WT IRE1α enabled complete RNA decay (Fig. 5b). Moreover, only WT IRE1α—but not IRE1α-R887A—depleted RIDDLE-targeted mRNAs TNFAIP8L1, SIX2, and SNN (Fig. 5b), We obtained similar data in HCC1806 cancer cells (Fig. S5b,c). Thus, IRE1α displays the same two basic endoribonuclease modalities in cells as does its KR module *in vitro*.

**Figure 5.**
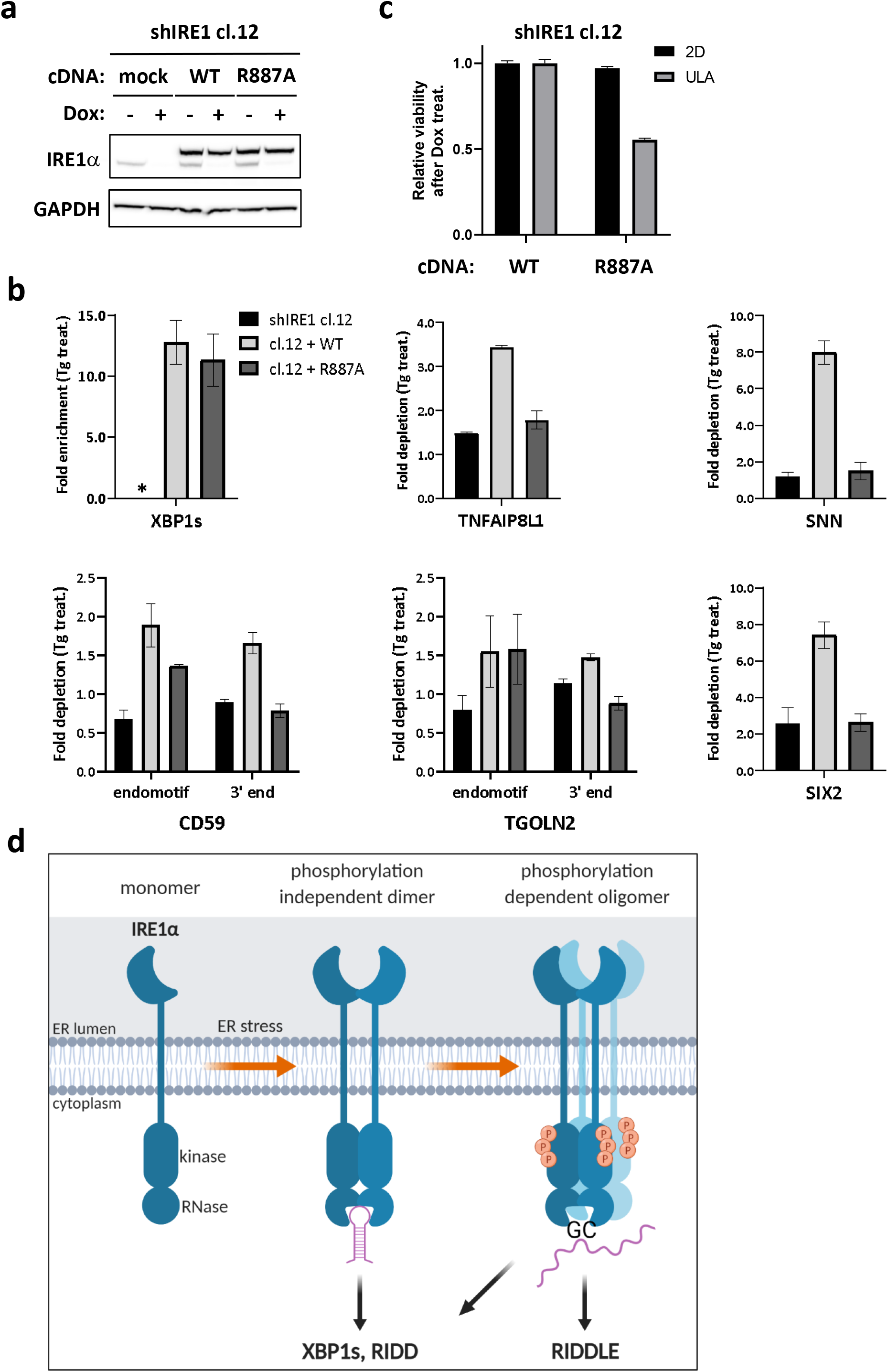
Cellular IRE1α displays the same fundamental RNase modalities and requires RIDDLE to support viability. (**a**) Western blot analysis of endogenous and ectopic IRE1α variant expression in MDA-MB-231 cells harboring Dox-inducible IRE1α shRNA stably transfected with transgenic WT or R887A mutant versions of IRE1α-GFP. (**b**) RT-qPCR analysis of IRE1α RNase targets CD59, TGOLN2 (RIDD), and TNFAIP8L1,SNN, and SIX2 (RIDDLE). *: Ct values for XBP1 in sample shIRE1 cl.1 prior to Tg treatment were >34, precluding ratio calculations. (**c**) Analysis of cell viability by Cell-Titer Glo after Dox treatment for 7 days on standard flat bottom (2-D) or Ultra-Low Attachment (ULA) plates (3-D growth). (**d**) Model depicting IRE1α’s principal modes of endoribonuclease function and their underlying phospho-oligomeric states during ER stress.

The failure of IRE1α-R887A to oligomerize and to acquire the corresponding RIDDLE modality afforded a unique opportunity to test the functional significance of the latter. To this end, we leveraged earlier work demonstrating that certain breast cancer cell lines depend on IRE1α for viability during 3D growth^14^. In both cell lines, transgenic WT IRE1α indeed rescued the loss of viability conferred by Dox-inducible knockdown of endogenous IRE1α; by contrast, IRE1α-R887A failed comparably to restore cell survival (Fig. 5c and Fig. S5d). Together, these results link IRE1α’s capacity to oligomerize and perform RIDDLE with its ability to support breast cancer cell survival during 3D growth.

## DISCUSSION

Our studies conceptually advance the current mechanistic understanding of IRE1α by shedding new light on the fascinating, yet puzzling, process of IRE1-dependent mRNA decay. Although in fly cells this process is thought to be relatively unrestricted by substrate sequence, in mammalian cells it was thought to require an XBP1-like stem-loop endomotif^40–44^. Despite some tantalizing clues that human IRE1α may degrade a wider scope of mRNAs^43,51^, to date this apparent disparity has not been deciphered. Our dual next-generation sequencing strategy, coupled with the discovery of a second basic endoribonuclease modality of IRE1α and its coordinate phospho-oligomeric state, enabled several advances: (1) a refinement of the canonical stem-loop endomotif; (2) the development of a powerful and accurate algorithm to discern such endomotifs in prospective mRNAs; and perhaps more importantly, (3) the identification of RIDDLE as a specific, biologically significant activity of human IRE1α.

As previously considered^40–42^, a majority of RIDD substrates did not harbor secretion signals, and nor did the newly uncovered RIDDLE targets. Given that IRE1 is an ER-resident membrane protein, these observations raise the fundamental question of how it gains access to mRNAs that are not translated by membrane-bound ribosomes. Future work will be required to decipher whether such target mRNAs are delivered in a non-conventional manner—independent of a signal sequence—to the ER, perhaps to sites of IRE1 cluster formation. Alternatively, a fraction of IRE1 molecules may become selectively proteolyzed to allow severed IRE1-KR domains to venture into the cytosol and act on non-membrane bound mRNAs^65,66^.

Transcripts decayed through both the RIDD and RIDDLE modalities showed enrichment in functional categories such as cell morphology and cell death or survival, which may be linked to the cellular response to ER stress. For example, DGAT2 promotes triglyceride synthesis^44^, while TNFAIP8L1 promotes apoptosis^67^.

Previous reasoning divided IRE1’s endoribonuclease modality into XBP1 processing *versus* RIDD. It was further thought that variation in target-sequence selectivity for RIDD substrates was probably due to inter-species diversity. Our findings establish a new conceptual organization (Fig. 5d) based on the two endoribonuclease modalities of human IRE1α described here. The first modality, which minimally requires IRE1α dimerization but can be performed also within phospho-oligomers (likely by the dimer-building blocks from which it is assembled), carries out endomotif-specific RNA cleavage; the second, which strictly requires phospho-oligomerization, conducts more promiscuous endoribonuclease activity. The first modality enables both the dual cleavage of XBP1u and initial single-site cleavage of RIDD targets containing a robust XBP1-like stem-loop endomotif. The second modality mediates RIDDLE, which digests RNA substrates that either have a sub-optimal stem-loop endomotif, or lack one altogether. Importantly, RIDDLE also degrades canonical endomotif-cleaved RIDD substrates. Our detailed studies with DGAT2 and TNFAIP18L1 RNA suggest that RIDDLE favors GC sites; future studies should examine additional RIDDLE substrates to determine whether such sites are universal. Our new conceptualization aligns IRE1α multimer assembly with endoribonuclease modality. Phosphorylated oligomers performed RNase cleavage with better efficiency than dimers, pointing to a “rheostatic” nature of IRE1α activation, as previously suggested^34,35^. Accordingly, stronger ER stress could drive higher levels of IRE1α phospho-oligomerization, increasing catalytic efficiency for both XBP1s generation and RNA decay.

Mutations that impaired phosphorylation and/or oligomerization confirmed and reinforced the requirement of distinct oligomeric states for the two basic RNase modes of IRE1α. Each phosphorylation site appeared similarly important for efficient oligomerization as well as for RNA decay, with triple phosphorylation leading to strongest activity. The data in breast cancer cells differs from B cells, wherein S729 proved more critical for RIDD^50^. Regardless, the R887A mutation, which prevented IRE1-KR oligomerization independent of phosphorylation state, demonstrated the critical importance of oligomers for RIDDLE-mediated RNA decay. Residing on human IRE1α B2B dimers, R887 may enable oligomer stabilization via double salt bridging with E876 residues on opposing B2B dimers. Supporting this notion, overlay of the human IRE1α dimer onto the yeast oligomer places E876 within just 7 Å from R887 (Fig. 4d).

RNase activity of IRE1α may vary with substrate length. It is possible that dual XBP1u processing occurs more efficiently within phosphor-IRE1α tetramers, which bind simultaneously to the two stem-loop endomotifs. Similarly, it is conceivable that for canonical RIDD targets, one IRE1α dimer binds to the stem-loop endomotif, while additional associated dimers within an oligomer cleave the transcript at additional locations. Future structural work is needed to explore oligomeric mammalian IRE1α alone and in complex with different RNAs. In addition, it will be important to investigate how the RNase modalities unraveled here apply to IRE1 from different species and model organisms. Emerging data for yeast IRE1 indicates that the positioning of the RNase domains within dimers regulates substrate recognition (W.L., P.W., submitted).

Intriguingly, even the initial products of XBP1u processing underwent detectable degradation by IRE1-KR-3P, consistent with the previously proposed concept that XBP1 mRNA splicing involves kinetic competition between exon ligation and mRNA degradation^51^. This earlier study, however, identified components of the NO-GO and cytosolic exosome machineries as the exonucleases mediating further decay from the primary cut sites. Importantly, although additional endonucleotic cleavages near ribosomes stalled on severed mRNA were uncovered, the nuclease(s) responsible for these cuts remained elusive. It remains to be determined whether IRE1 plays a direct part in such clearing mechanisms by which cells rid themselves of defective mRNA. In the cellular environment, additional factors, including the Sec61 translocon^68^, or RNA-binding proteins such as Pumilio (Cairrao et al, co-submitted), could also help shield processed XBP1u against further decay and thereby permit exon ligation by RtcB^69^.

In conclusion, our discovery and functional characterization of two enzymatic modalities of IRE1α RNase advance the present conceptual framework for investigating how IRE1 operates across different eukaryotes. Together with the demonstration that RIDDLE is important for cancer-cell survival, our mechanistic dissection carries important implications for biological understanding of the UPR and for harnessing IRE1α as a potential therapeutic target.

## Supporting information

gRIDD Supplemental Methods

## ACKNOWLEDGEMENTS

We thank Ben Haley for gDNA design, Maria Lorenzo for protein purification, Maureen Beresini, Ira Mellman, and members of the Ashkenazi lab at Genentech and the Walter lab at UCSF for helpful discussions.

## FIGURE LEGENDS

**Figure S1.**
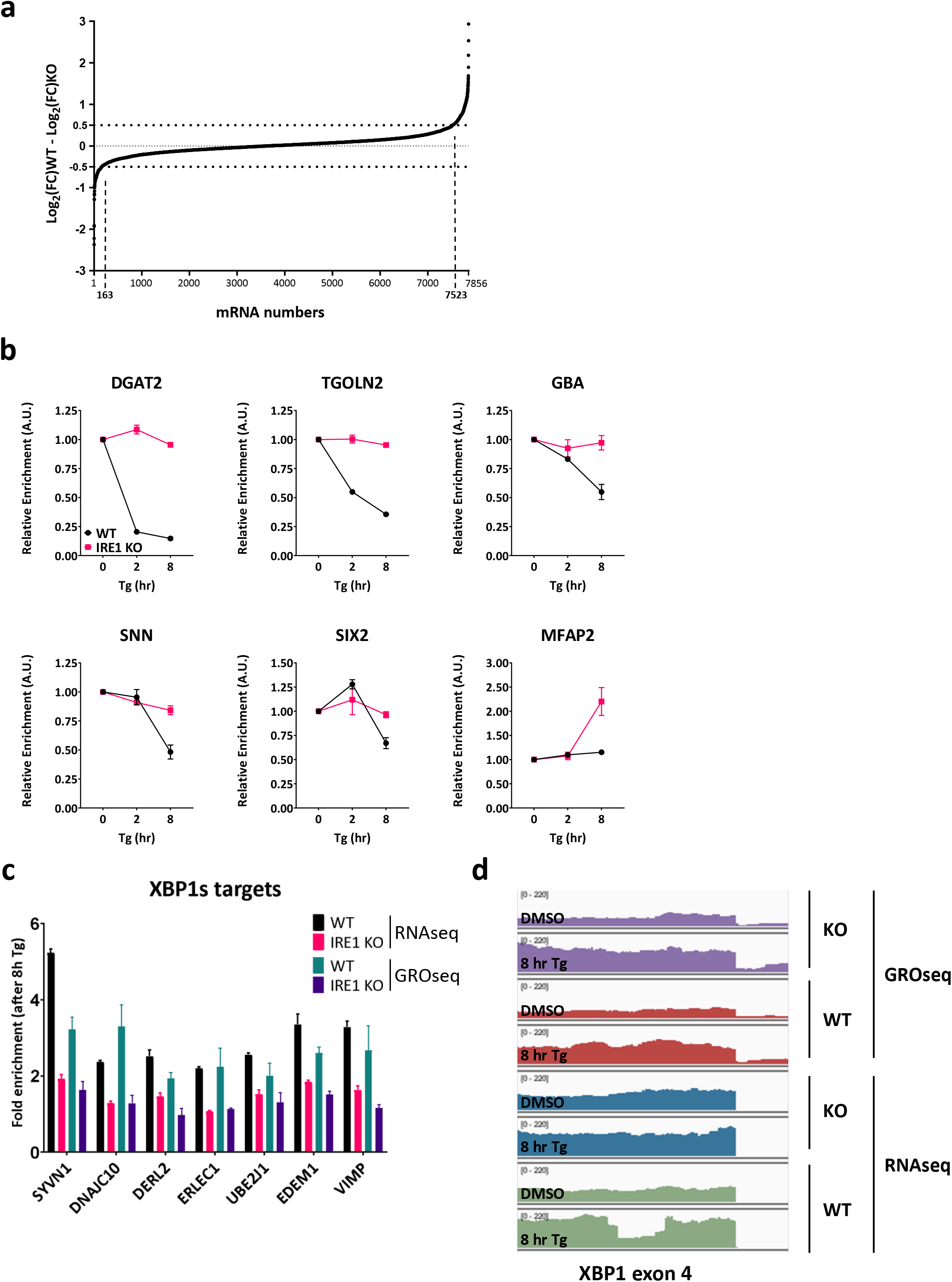
Integrative RNAseq and GROseq analyses identify human RIDD and RIDDLE targets. (**a**) IRE1-dependent Log_2_ fold change in mRNA levels after Tg treatment of MDA-MB-231 cells, as determined by RNAseq. (**b**) Kinetic RT-qPCRs analysis of gene transcripts identified through integrative RNAseq and GROseq analysis in *IRE1α* WT and KO MDA-MB-231 cells. (**c**) Fold enrichment after ER stress induction by Tg for specific XBP1s target genes in IRE1α WT and KO cells, as determined from the RNAseq and GROseq datasets. (**d**) Read coverage around the spliced XBP1 genomic region (part of exon 4) in all various datasets.

**Table S1.** Summary of IRE1 dependent down-regulated targets in MDA-MB-231 cells. mRNA transcripts down-regulated upon ER stress in an IRE1α-dependent and transcription-independent manner. The percentage of IRE1α-dependent down-regulation upon Tg treatment measured by RNAseq is represented in column 3. Fold depletion of total RNA (RNAseq Log_2_(FC)WT) over transcriptional down-regulation (GROseq Log_2_(FC)WT) is indicated in column 4. Analysis for the presence of a signal peptide (Type I) or signal anchor (Type II) sequence is indicated in the last column.

**Figure S1**
|  | Gene name | IRE1 dep. down-reg.<br>(RNAseq) | Fold down-reg. over<br>transcription<br>(RNAseq/GROseq) | Signal<br>sequence |
| --- | --- | --- | --- | --- |
| 1 | AIM2 | 37% | 1.67 | None |
| 2 | ALDH1A3 | 37% | 1.25 | None |
| 3 | ATF5 | 35% | 1.58 | None |
| 4 | ATP9A | 41% | 1.44 | None |
| 5 | B3GNT8 | 39% | 1.89 | Type I |
| 6 | BCAM | 79% | 1.71 | Type I |
| 7 | BLOC1S1 | 30% | 1.31 | None |
| 8 | BMP4 | 49% | 1.32 | Type I |
| 9 | CALHM2 | 31% | 1.33 | None |
| 10 | CCDC69 | 33% | 1.41 | None |
| 11 | CD59 | 42% | 1.65 | Type I |
| 12 | CDKN1C | 47% | 1.65 | Type I |
| 13 | CEP162 | 41% | 1.93 | None |
| 14 | CFAP45 | 32% | 1.96 | None |
| 15 | CHST4 | 45% | 3.29 | None |
| 16 | CTSO | 45% | 1.33 | Type I |
| 17 | DGAT2 | 81% | 4.07 | Type II |
| 18 | DSEL | 37% | 1.30 | Type I |
| 19 | ERCC6 | 30% | 1.28 | None |
| 20 | FAM117B | 30% | 1.43 | None |
| 21 | FAM63A | 53% | 1.29 | None |
| 22 | FILIP1L | 50% | 1.31 | None |
| 23 | FITM2 | 31% | 2.18 | Type II |
| 24 | GATSL2 | 29% | 1.51 | None |
| 25 | GBA | 41% | 1.38 | Type I |
| 26 | GJD3 | 32% | 1.36 | Type I |
| 27 | GPC1 | 33% | 1.34 | Type I |
| 28 | HAPLN3 | 40% | 2.01 | Type I |
| 29 | HIP1 | 40% | 1.45 | None |
| 30 | IL31RA | 41% | 1.27 | Type I |
| 31 | KIAA1467 | 33% | 1.58 | None |
| 32 | LRRN4 | 32% | 3.05 | Type I |
| 33 | MAP3K5 | 31% | 1.30 | None |
| 34 | MAST4 | 35% | 1.30 | None |
| 35 | METTL7A | 41% | 2.15 | None |
| 36 | MFAP2 | 37% | 1.90 | Type I |
| 37 | MILR1 | 42% | 2.12 | Type I |
| 38 | OAS2 | 41% | 1.32 | None |
| 39 | PIGQ | 33% | 1.38 | None |
| 40 | PROS1 | 38% | 4.38 | Type I |
| 41 | RGS7 | 35% | 1.39 | None |
| 42 | RNF213 | 33% | 1.41 | None |
| 43 | SIX2 | 37% | 1.60 | None |
| 44 | SNN | 32% | 1.78 | Type II |
| 45 | SRXN1 | 36% | 1.24 | None |
| 46 | SUOX | 32% | 1.32 | None |
| 47 | TGOLN2 | 56% | 2.43 | Type I |
| 48 | TLR2 | 43% | 1.68 | Type I |
| 49 | TMEM19 | 37% | 1.86 | None |
| 50 | TNFAIP8L1 | 30% | 4.04 | None |
| 51 | TRIM16L | 32% | 1.51 | None |
| 52 | TRIM62 | 30% | 1.34 | None |
| 53 | WT1 | 50% | 1.49 | None |
| 54 | ZFHX4 | 36% | 1.27 | None |

**Table S2.** Ingenuity Pathway Analysis (IPA) of IRE1α’s mRNA decay targets in MDA-MB-231 cells. Top 5 enriched pathways in the Molecular and Cellular Functions category according to the IPA software, along with their p-value range and the number of targets in that group.

| **Molecular and cellular functions** | **p-value range** | **# genes** |
| --- | --- | --- |
| Cell death and survival | 1.41e-02 – 6.16e-05 | 11 |
| Cell signaling | 1.18e-02 – 7.77e-05 | 4 |
| Post-translational modification | 8.91e-03 – 7.77e-05 | 6 |
| Cell morphology | 1.41e-02 - 2.44e-04 | 10 |
| Cell cycle | 1.41e-02 - 3.75e-04 | 8 |

**Table S3.** GO term analysis of mRNA targets up-regulated in IRE1α-dependent manner after Tg treatment in MDA-MB-231 cells based on the GROseq dataset. Most enriched GO terms in the Biological process category, along with their p-value and the number of targets in that group.

| **GO biological process** | **p-value** | **# genes** |
| --- | --- | --- |
| Response to endoplasmic reticulum stress (GO:0034976) | 1.87E-31 | 46 |
| Endoplasmic reticulum to Golgi vesicle-mediated transport (GO:0006888) | 2.89E-26 | 37 |
| IRE1-mediated unfolded protein response (GO:0036498) | 6.73E-17 | 18 |
| Cellular response to unfolded protein (GO:0034620) | 3.78E-16 | 23 |
| Protein N-linked glycosylation via asparagine (GO:0018279) | 3.80E-15 | 14 |
| ERAD pathway (GO:0036503) | 8.90E-15 | 19 |
| COPII-coated vesicle cargo loading (GO:0090110) | 1.46E-12 | 10 |
| Protein folding in endoplasmic reticulum (GO:0034975) | 3.74E-06 | 5 |
| Somatostatin signaling pathway (GO:0038170) | 7.45E-06 | 4 |
| SRP-dependent cotranslational protein targeting to membrane (GO:0006614) | 7.50E-06 | 10 |
| COPI coating of Golgi vesicle (GO:0048205) | 1.23E-05 | 4 |
| Protein K69-linked ufmylation (GO:1990592) | 2.11E-04 | 3 |
| Mannose trimming involved in glycoprotein ERAD pathway (GO:1904382) | 2.11E-04 | 3 |

**Figure S2.**
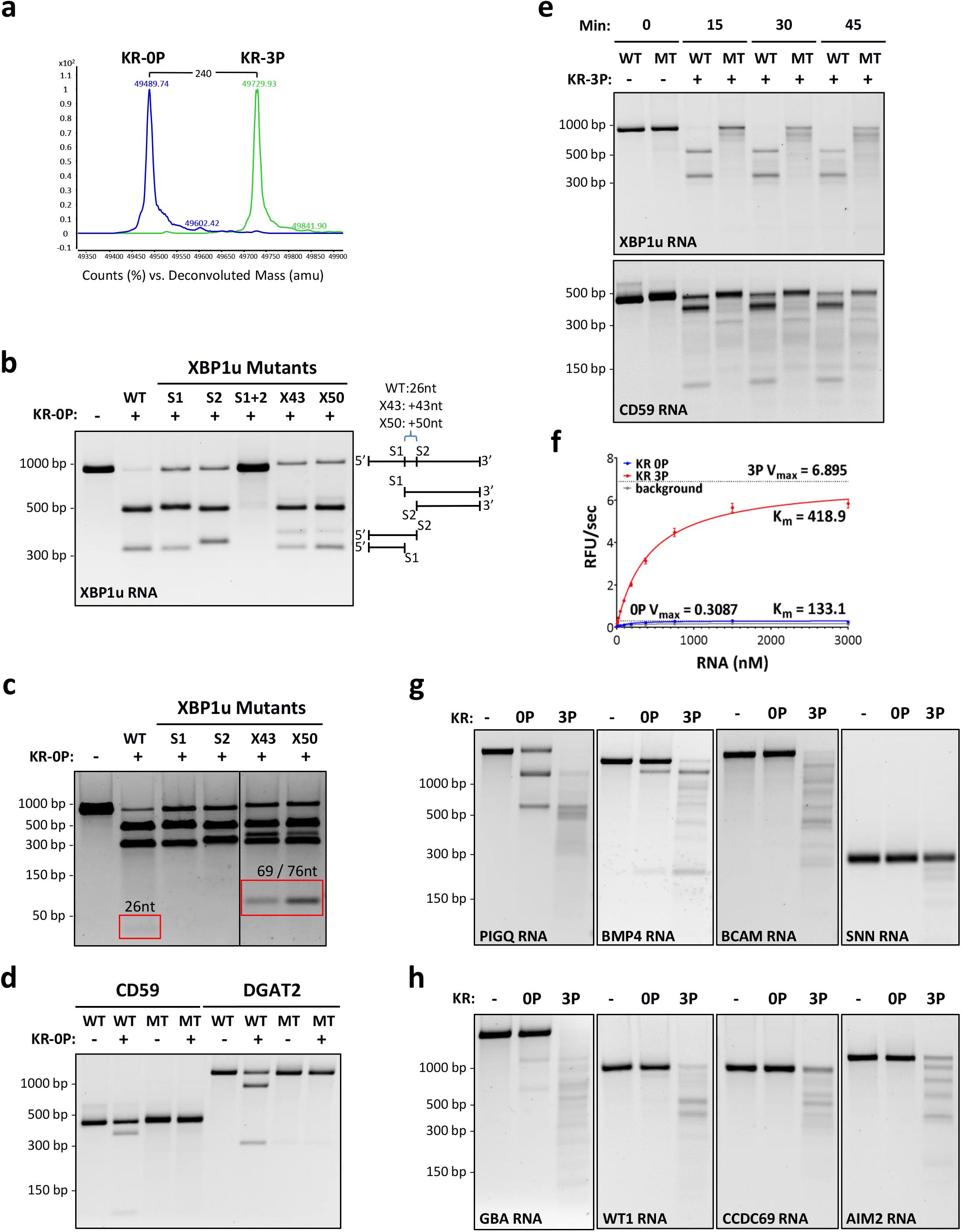
Phosphorylation state of IRE1α affects RNase modality. (**a**) Each IRE1-KR protein was purified by size exclusion chromatography and its identity and phosphorylation state were verified by liquid chromatography-mass spectrometry (LC-MS). Pre-existing phosphates were removed by treatment with λ-phosphatase (KR-0P), or full autophosphorylation was allowed in the presence of ATP (KR-3P). (**b**) KR-0P digestion of XBP1u transcript variants: WT; loop motif 1 scrambled (S1); loop motif 2 scrambled (S2); loop motif 1 and 2 scrambled (S1+2); intron with 43-nt (X43) or 50-nt (X50) random sequence inserted. Schematic illustration of each variant is depicted on the right side of the gel aligned with corresponding products. (**c**) KR-0P digestion of XBP1u variants at higher concentrations to visualize resulting spliced fragments (highlighted in the red boxes with their respective expected size). (**d**) KR-0P digestions of CD59 and DGAT2 RNA transcripts in WT and scrambled endomotif mutant (MT) version. (**e**) KR-3P digestions of WT or MT (scrambled endomotif) versions of XBP1u, DGAT2, and CD59 RNAs were performed for the indicated time and analyzed as above. (**f**) Michaelis-Menten kinetics for RNase activity of KR 0P (blue) and KR-3P (red). Each KR protein (10 nM) was incubated for 1 hr at room temperature with quenched fluorescein-conjugated RNA substrate at varying concentrations. As the substrate is cleaved by IRE1α RNase, FAM fluorescence is emitted and can be measured at regular intervals during the incubation time (see methods section for details). Velocity was measured as Relative Fluorescent Units (RFU)/sec and is shown in the graph as a function of RNA substrate concentration. Background signal of the RNA-only sample is depicted in gray. Kinetic parameters (Km and Vmax) of the reaction were calculated for both enzymes using Prism and are reported on the graph. (**g**,**h**) T7 RNAs for PIGQ, BMP4, BCAM, and SNN (**g**); or GBA, WT1, CCDC69, and AIM2 (**f**) were incubated with KR-0P and KR-3P and analyzed.

**Table S4.**
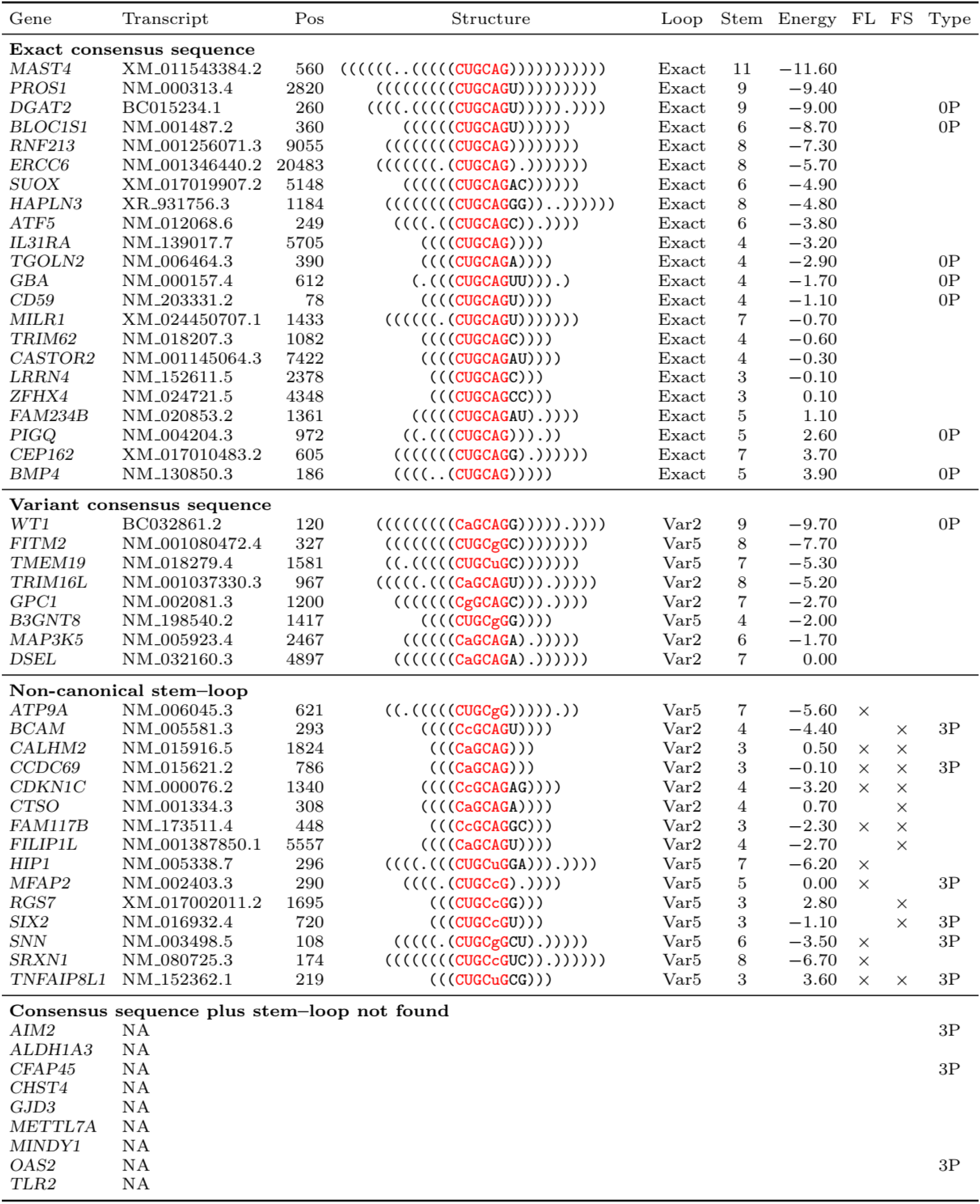
Canonical stem-loop endomotif analysis of RIDD targets from Table S1 using the gRIDD algorithm. mRNA transcripts down-regulated upon ER stress in an IRE1α-dependent and transcription-independent manner were analyzed for the presence and robustness of a canonical XBP1-like stem-loop endomotif, using a newly developed algorithm termed gRIDD (see supplementary methods for detailed description of gRIDD). Columns 1 and 2 indicate gene names and transcript references. Column 3 indicates mRNA nucleotide position of the G within the GC cleavage site. Column 4 illustrates the loop sequence and stem topology of the endomotif, where a parenthesis indicates a nucleotide that is paired with a complementary one on the opposite leg of the stem, and a dot indicates an unpaired nucleotide. Lowercase letters indicate variation to the consensus loop sequence at position 2 or 5, indicated in column 5 as Var2 and Var5 respectively. Column 6 indicates the number of base pairs at the stem. Column 7 indicates the free energy (kcal/mol) for the stem-loop structure represented in column 4. Columns 8 and 9 indicate failure to meet gRIDD criteria due to excessive loop-length (FL) and/or disrupted base pairing at the stem (FS). Column 10 indicates empirical results for cleavage by IRE1-KR-0P (0P) or IRE1-KR-3P (3P). All transcripts cleaved by IRE1-KR-0P were also cleaved by IRE1-KR-3P. In each case, the best possible stem-loop endomotif is displayed. The first and second categories include mRNAs with stem-loop endomotifs that meet gRIDD criteria (RIDD modality). The third and fourth categories include mRNAs wherein the best possible endomotif nevertheless fails to meet gRIDD criteria or no stem-loop endomotif is found (RIDDLE modality). Within each category, transcripts are ranked by free energy.

**Figure S3.**
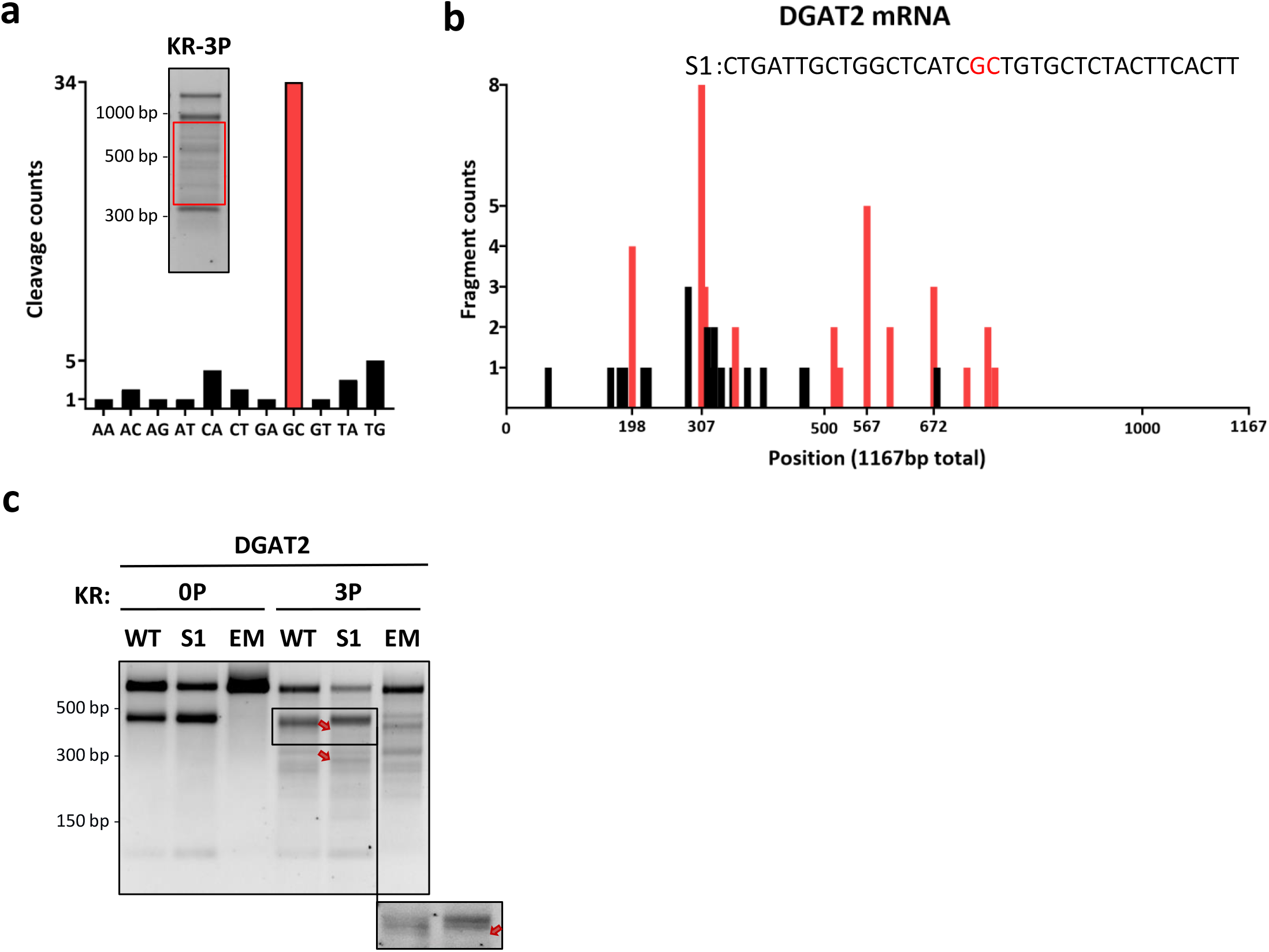
RIDDLE is relatively unrestricted but non-random. (**a**) Amount of RNA fragments sequenced whose 3’ end leads to the cleaved nt pair designated on the x axis. The first nucleotide in the pair represents the last sequenced nucleotide from the RNA fragment, while the second nucleotide shows the following base in the RNA sequence. Inset: red box indicates the portion of the gel that was extracted for subsequent sequencing. (**b**) Mapping of the last base pair (3’ end) from each individual RNA fragment sequenced within the DGAT2 mRNA. Red bars indicate cleavage sites between a GC nt pair. (**c**) RNA digestions of WT DGAT2 and DGAT2 mutated at location S1 or at the stem-loop endomotif (EM). The red arrows indicate change in banding pattern as compared to WT.

**Figure S4.**
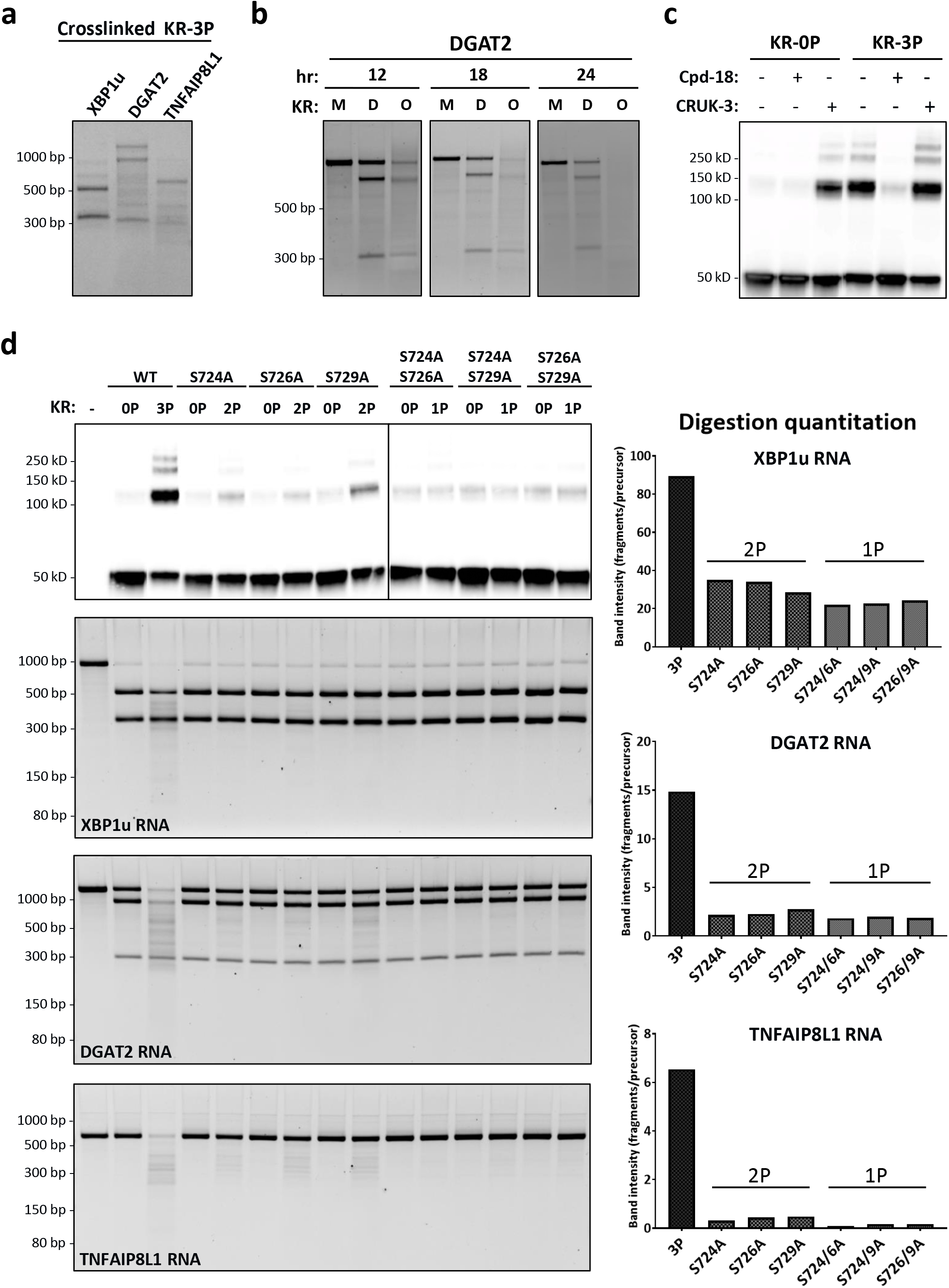
Phospho-oligomeric state governs IRE1α’s RNase modality. (**a**) IRE1-KR-3P digestion of XBP1u, DGAT2, and TNFAIP8L1 RNA immediately after DSS crosslinking and before fractionation of the protein. (**b**) Digestion of DGAT2 RNA by crosslinked IRE1-KR-3P at the indicated incubation time after protein fractionation. (**c**) DSS crosslinking of IRE1-KR in the presence of Cpd-18 or CRUK-3. (**d**) DSS crosslinking of IRE1-KR WT or IRE1-KR mutated at the indicated activation-loop phosphorylation sites, and corresponding digestion of XBP1u, DGAT2, and TNFAIP8L1 RNA. Intensities of RNA fragments for each variant were quantified using GelQuantNET, as depicted on the right side of the gel.

**Figure S5.**
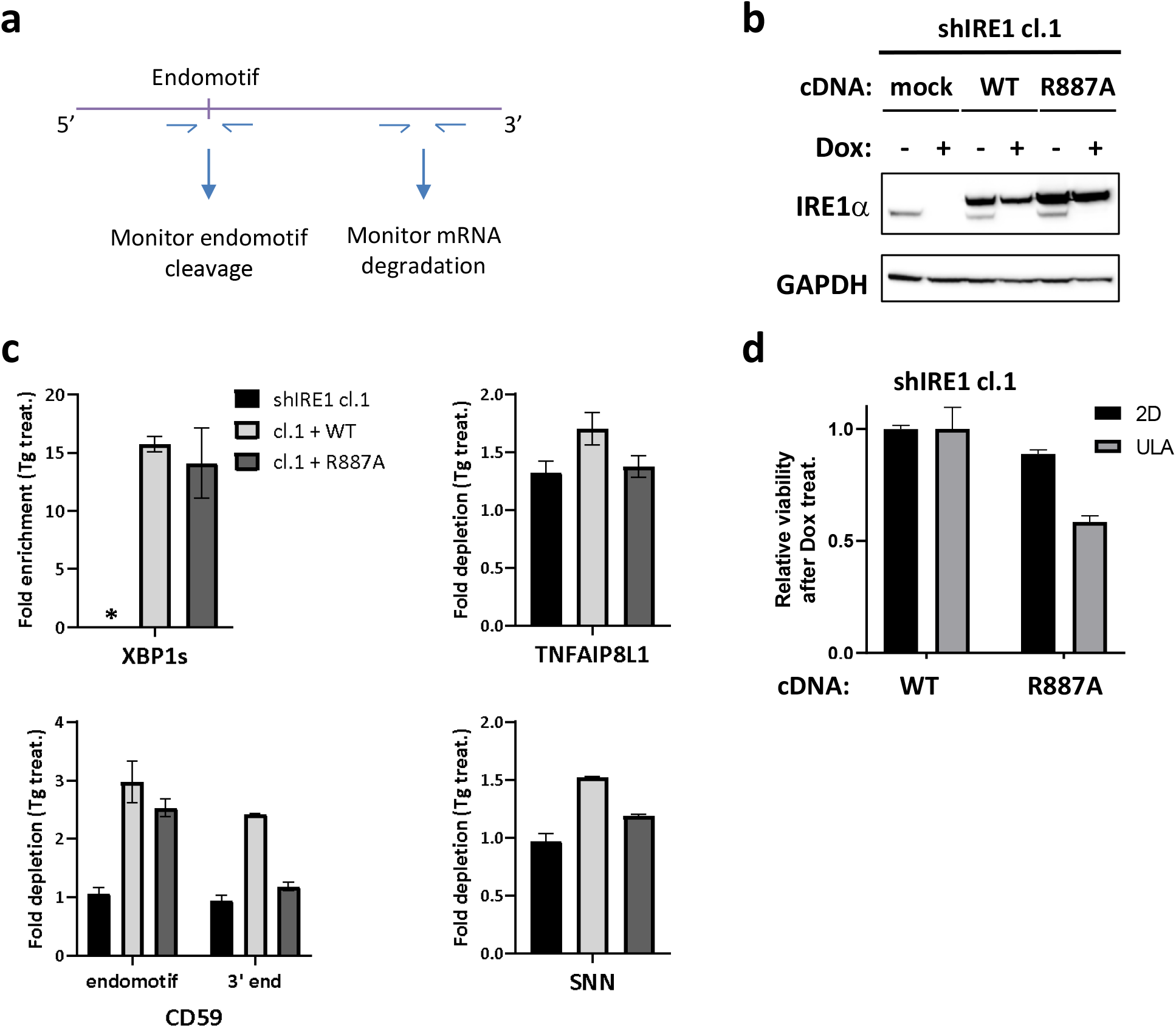
Cellular IRE1α displays the same two fundamental RNase modalities and requires RIDDLE to support viability. (**a**) Cartoon depicting the location of the primer pairs used for RT-qPCR analysis of endomotif-directed cleavage vs. RIDDLE. (**b**) Western blot analysis of endogenous and ectopic IRE1α variant expression in HCC1806 cells harboring Dox-inducible IRE1α shRNA that were stably transfected with transgenic WT or R887A mutant versions of IRE1α-GFP. (**b**) RT-qPCR analysis of IRE1α RNase targets as described for Fig. 5.

## MATERIALS AND METHODS

### Cell culture and experimental reagents

MDA-MB-231 cells were all obtained from ATCC, authenticated by short tandem repeat (STR) profiles, and tested to ensure mycoplasma free within 3 months of use. All cell lines were cultured in RPMI1640 media supplemented with 10% (v/v) fetal bovine serum (FBS, Sigma), 2 mM glutaMAX (Gibco) and 100 U/ml penicillin plus 100 μg/ml streptomycin (Gibco).

Thapsigargin (Sigma) was used at a concentration of 100 nM. Compound 4µ8C and Compound 18 were dissolved in DMSO for cellular experiments and used at the indicated concentrations. Antibodies (Abs) for IRE1α (#3294), and GAPDH (#8884) are from Cell Signaling Technology. Secondary antibodies (rabbit #711-035-152 and mouse #715-035-150) were from Jackson ImmunoResearch Laboratories.

### CRISPR/Cas9 knockout: Guide RNA sequences and technique

MDA-MB-231 *IRE1*α KO cells were generated using CRISPR by co-transfecting a Cas9 containing plasmid, pRK-TK-Neo-Cas9, with a pair of IRE1 targeting gRNAs (see below) cloned into a pLKO vector. Transfection was done using Lipofectamine 3000 according to manufacturer protocol, and transformants were selected by PCR on genomic DNA for the detection of deletions. Correct clones were then sequenced. *IRE1*α KO cl.1-2 gRNA pair: CTTGTTGTTTGTGTCAACGC & TCTTGCTTCCAAGCGTATAC.

### RNAseq / GROseq

Both RNAseq and GROseq were performed on WT and *IRE1*α KO MDA-MB-231 cells. For each RNAseq and GROseq there were four experimental conditions (WT and *IRE1*α KO, treated with Tg or vehicle control (DMSO) for 8 hr with three biological replicates (n=3) for each condition.

For RNAseq, RNA was extracted using the RNeasy kit (Qiagen #74104) performing on-column DNA digestion for 15 min. The concentration of RNA samples was determined using NanoDrop 8000 (Thermo Scientific) and the integrity of RNA was analyzed by Fragment Analyzer (Advanced Analytical Technologies). Approximately 500 ng of total RNA was used as input for library preparation using TruSeq RNA Sample Preparation Kit v2 (Illumina).

For GROseq, cells were pre-treated with 5-ethynyl uridine (EU) for 30 min prior to nuclei fractionation using Sigma kit (#NUC101) followed by Invitrogen Click-iT™ Nascent RNA Capture Kit protocol for nascent transcript extraction (#C10365). RNA was then extracted using the RNeasy kit (Qiagen #74104) performing on-column DNA digestion for 15min and was subsequently reverse transcribed with Superscript VILO IV Master Mix from Invitrogen. Libraries were prepared following the protocol from NEBNext® Ultra™ II RNA Library Prep Kit for Illumina® (NEB #E7770S) and indexes (NEB #E7335S and #E7500S).

The size of the libraries for both RNAseq and GROseq was confirmed using 4200 TapeStation and High Sensitivity D1K screen tape (Agilent Technologies) and their concentration was determined by qPCR based method using Library quantification kit (KAPA). The libraries were multiplexed and then sequenced on Illumina HiSeq2500 (Illumina) to generate 30M of single end 50 base pair reads.

### RNAseq-GROseq combined analysis of IRE1α specific targets

To identify IRE1α specific RIDD targets we compared genes’ differential expression in WT *versus* KO cells after 8 h Tg treatment: Log_2_ fold change (Log_2_(FC)), average expression, *p*-values, and false discovery rate (fdr) values were calculated for every protein-coding genes comparing the starting DMSO time point with 8 h after Tg treatment in WT and *IRE1*α KO conditions for both RNAseq and GROseq datasets. Log_2_(FC) in the RNAseq WT dataset for which the *p*-value and the fdr value were above 0.05 were removed. Log_2_(FC) difference between the RNAseq WT and *IRE1*α KO (Log_2_(KO-WT)) datasets falling below 0.5 were removed, and genes with average expression values below 1 were also removed. Finally, Log_2_(FC) differences between the RNAseq and GROseq WT (Log_2_(WTrna-WTgro)) falling below −0.3 were removed. The resulting gene list is 55 entries long.

### Signal sequence analysis

For each mRNA transcript selected as the representative candidate for a gene, we determined the corresponding protein sequence by running GMAP (version 2019-12-01)^70^ by aligning the sequence to itself using the -g flag and extracting the full-length protein translation with the flags “-P -F”. We then ran the program signalp (version 3.0)^71^ on the protein sequence with the flag “-t euk”, which yielded a signal sequence prediction and probabilities for the signal peptide, signal anchor, and cleavage site. Each mRNA was also checked in Uniprot for additional verification.

### RT-qPCRs

RNA was extracted using the RNeasy Plus kit (Qiagen #74134). Equal amounts of RNA were reverse transcribed and amplified using the TaqMan™ RNA-to-CT™ 1-Step Kit (Applied Biosystems #4392938) on the ABI QuantStudio 7 Flex Real-Time PCR System. The delta-delta C_T_ values were calculated by relating each individual C_T_ value to its internal GAPDH control. Taqman primers for XBP1u (#Hs02856596_m1), XBP1s (#Hs03929085_g1), DGAT2 (#Hs01045913_m1), BLOC1S1 (#Hs00155241_m1), CD59 (#Hs00174141_m1), and TNFAIP8L1 (#Hs00537038_m1), and GAPDH (#Hs02758991_g1) were from Life Technology. Additional primer pairs used for qPCR from cell rescue experiments were ordered from IDT: TNFAIP8L1 (#Hs.PT.58.39992641), SNN (#Hs.PT.58.28146300), SIX2 (#Hs.PT.58.40614621), GAPDH (#Hs.PT.39a.22214836), and custom-designed:

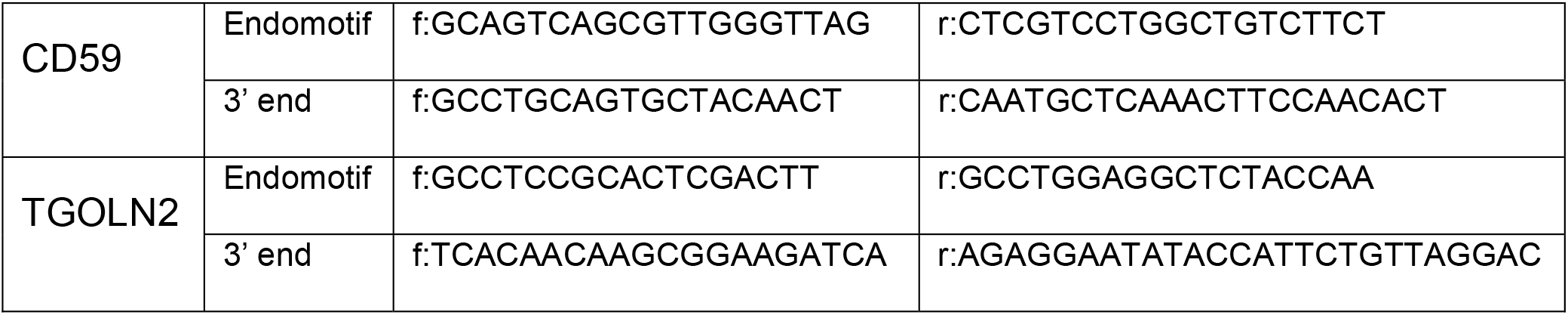

### T7 RNA constructs

We prepared T7 RNA transcripts from cDNA templates chosen based upon functional relevance coupled with optimal length for the ribonucleolytic reaction (∼0.5 to 2 kb). cDNA constructs encoding XBP1 (#HG10751-UT), DGAT2 (#HG14114-G), CD59 (#HG12474-UT), TGOLN2 (#HG17252-UT), SIX2 (#HG21116-UT), CFAP45 (#HG22377-UT), MFAP2 (#HG16644-UT), PIGQ (#HG22757-UT), BMP4 (#HG10609-UT), BCAM (#HG10238-UT), SNN (#HG23279-U), GBA (#HG12038-UT), WT1 (#HG12282-UT), CCDC69 (#HG27177-U), AIM2(#HG11654-UT) were from Sino Biological, and BLOC1S1 (#RC224412), TNFAIP8L1 (#RC203912) from Origene. cDNA was amplified using T7 forward primers, and subsequently in-vitro transcribed using HiScribe™ T7 Quick High Yield RNA Synthesis Kit from NEB (#E2050S).

T7 RNA mutations were engineered using overlap PCR followed by restriction digests, and the final fragments were purified from agarose gel (Zymoclean Gel DNA Recovery kit #D4001).

### Protein purification and separation of phosphorylated IRE1α fractions

IRE1α KR 0P and 3P were produced by Accelagen and in-house: IRE1α KR (G547-L977) was expressed as N-terminal His_6_-tagged fusion proteins in SF9 cells with a TEV protease cleavage site from an intracellular BEVS expression vector. Cell pellet was resuspended in lysis buffer containing 50 mM HEPES pH 8.0, 300 mM NaCl, 10% glycerol, 1 mM MgCl2, 1:1000 benzonase, EDTA-free PI tablets (Roche), 1mM TCEP, and 5mM imidazole. Sample was lysed by sonication, centrifuged at 14,000rpm for 45min, and the supernatant filtered through a 0.8 μm□Nalgene filter. Cleared supernatant was bound to Ni-NTA Superflow beads (Qiagen) by gravity filtration. Beads were washed in lysis buffer supplemented with 15 mM imidazole, followed by protein elution in lysis buffer containing 300 mM imidazole. The eluate was incubated with TEV protease overnight at 4°C. The sample of IRE1α KR protein was diluted 1:10 in 50 mM HEPES pH 7.5, 50 mM NaCl, 1mM TCEP and then loaded onto a 5mL pre-packed Q-HP column (GE-Healthcare). Separation of IRE1α KR unphosphorylated and phosphorylated was achieved by eluting the protein with a very shallow gradient (50-300 mM NaCl over 70CV). Fully phosphorylated fraction (MW+240 by LC-MS) was collected separately, while the rest of the protein fractions were consolidated and incubated with Lambda phosphatase for one hour at room temperature.

Dephosphorylation was confirmed by LC-MS. Unphosphorylated and phosphorylated samples were then concentrated and loaded separately onto a HiLoad 16/600 Superdex 200 SEC column (GE Healthcare) equilibrated in 25 mM HEPES pH 7.5, 250 mM NaCl, 1 mM TCEP, 10% glycerol. IRE1α eluted as a monomer.

Mutants S724A, S726A, S729A, S724A-S726A, S724A-S729A, S726A-S729A, and R887A were produced in-house following the same procedure described above.

Phosphorylation site mapping was performed by LC-MS/MS analysis following protease digestion:

| **IRE1α KR** | **% phosphorylation** |  |  |
| --- | --- | --- | --- |
|  | **S724** | **S726** | **S729** |
| **WT 3P** | 82 | Yes* | 52 |
| **S724A 2P** | - | >90 | >90 |
| **S726A 2P** | 90 | - | 71 |
| **S729A 2P** | 82 | Yes* | - |
| **S724A-S726A 1P** | - | - | 97 |
| **S726A-S729A 1P** | 89 | - | - |
| **S724A-S729A 1P** | - | Yes* | - |

Yes\* denotes observed S726 phosphorylation for which quantification was not possible due to the presence of a miscleaved peptide containing phosphorylated S726.

### Phosphorylation of IRE1α KR and activation loop mutants

IRE1α KR S/A and R887A mutants were allowed to autophosphorylate in the presence of 2 mM ATP and 10 mM MgCl_2_ for one hour at room temperature. Sample was purified from ADP by size-exclusion chromatography (SEC).

Mutants S726A-S729A, was not capable to autophosphorylate and was instead incubated with pIRE1α LKR (Linker-Kinase-RNase, residues Q470-L977) at 1:40 w/w, 2 mM ATP and 20 mM MgCl_2_. Final phosphorylated proteins were purified from pIRE1α LKR and residual nucleotides by SEC.

### RNA cleavage assay

1μg of T7 RNA generated was digested at room temperature by 1 μg of human IRE1α KR recombinant protein (∼0.8 μM) for 15 min in RNA cleavage buffer (HEPES pH7.5 20 mM; K acetate 50 mM; Mg acetate 1 mM; TritonX-100 0.05% (v/v)). The total volume of the reaction is 25 μl. The digestion was then complemented by an equal volume of formamide and heated up at 70°C for 10 min to linearize the RNA. After linearization, the mixture was immediately placed on ice for 5 min, and then 20 μl was run on 3% agarose gel at 160V for 1 hour at 4°C. If inhibitors were used, they were incubated with the RNA for 40 min on ice prior to RNA digestion. Gels were visualized on either a BioRad Molecular Imager ChemiDoc ZRS+ or Azure Biosystems c600.

### RNA fragment sequencing

2 μg of T7 RNA (TNFAIP8L1, DGAT2) is used for digestion by human IRE1α KR-3P as described above. RNA bands are extracted from gel using Zymoclean Gel RNA Recovery Kit (Zymo Research #R1011). Next, the RNA extracted was ligated using RtcB ligase (NEB # M0458S) to a 3’-adapter oligo custom designed and ordered from IDT (caagcagaagacggcatacgagatCGTGAT), following manual protocol. Ligated RNA was then reverse transcribed using SuperScript™ IV First-Strand Synthesis System (Invitrogen #18091050) and a 3’-adapter specific primer (ATCACGatctcgtatgccg). cDNA was then amplified using DGAT2 (gggGCATGCATGAAGACCCTCATAGCCG) or TNFAIP8L1 (gggGCATGCATGGACACCTTCAGCACCAAG) specific forward primer containing an SphI restriction enzyme site, and a common reverse primer containing a EcoRI restriction enzyme site (gggGAATTCATCACGatctcgtatgccg). PCR product were subsequently disgested by SphI and EcoRI prior cloning into a pGEM®-T Easy Vector (Promega #A1360). Then, resulting plasmids are transfected into competent cells (Zymo *Mix & Go!* Competent Cells Zymo 10B #T3019), plated on a 10 cm dish and left at 37°C overnight. The following day, individual colonies are picked and grown onto 96 wells plates, designed for bacterial growth (Thomson Instrument Company #951657), overnight. Finally, DNA is extracted from individual wells using Zyppy-96 Well Plasmid Miniprep Kit (Zymo Research #D4042) and sent for SANGER sequencing.

### RNase activity assay (kinetic fluorescence)

A 5′-Carboxyfluorescein (FAM)-and 3′-Black Hole Quencher (BHQ)-labeled single stem-loop mini-substrate containing XBP1 sequence (5′FAM-CAUGUCCGCAGCGCAUG-3′BHQ) was used as substrate for cleavage by IRE1α KR (G547-L977) in 20 mM HEPES pH 7.5, 50 mM potassium acetate, 1 mM magnesium acetate, 1 mM dithiothreitol, 0.05% v/v TritonX-100. 10 nM of protein was incubated with varying concentrations of RNA substrate (2-fold dilution series from 3000 nM to 5.86 nM). RNA cleavage was measured kinetically over an hour at room temperature as an increase in fluorescence. The final reaction was carried out in 20 uL in 384-well plates. Samples were run in duplicate. Velocity of the reaction was measured as the slope of the linearly increasing fluorescence signal over time as Relative Fluorescence Units (RFU)/sec and plotted as a function of RNA substrate concentration. Michaelis-Menten kinetics were fit using Prism 1.7 and resulting Vmax and Km constant were reported.

### Immunoblot analysis

Cells were lysed in 1x RIPA buffer (Millipore) supplemented with fresh protease and phosphatase inhibitors (Invitrogen #78440), cleared by centrifugation at 13,000 rpm for 15 min, and analyzed by BCA protein assay (Thermofisher Scientific #23227). Equal protein amounts were loaded, separated by SDS-PAGE, electrotransferred to nitrocellulose membranes using the iBLOT2 system (Invitrogen), and blocked in 5% nonfat milk solution for 30 min. Membranes were probed with the following antibodies: IRE1α, XBP1s, or GAPDH. Signal was detected using appropriate horseradish peroxidase (HRP)-conjugated secondary antibodies. All primary antibodies were used at 1:000 dilution and overnight hybridization at 4°C, followed by a two-hour incubation with horseradish peroxidase (HRP)-conjugated secondary antibodies at 1:10,000 dilution.

### Crosslinking Assay

1 μg of IRE1-KR recombinant protein was crosslinked in 25 μl final volume of RNA cleavage buffer containing 1 μl of disuccinimidyl suberate (DSS, Thermo Fisher Scientific) crosslinker at 6.25 mM (final concentration is 250 μM) for 1 h on ice. For experiments recurring inhibitor and activator compounds, IRE1α KR and the small molecule compound were first pre-incubated together on ice for 40min. The reaction was quenched using 1 μl of a pH 7.5, 1M TRIS solution for 15 min on ice. The reaction was then diluted in 500 μl of RNA cleavage buffer and 13 μl of it was used to run on SDS-PAGE gel (corresponding to ∼25 μg of protein. The gel was run at 100V for almost 3 h then electro-transferred to nitrocellulose membranes using the iBLOT2 system (Invitrogen), and blocked in 5% nonfat milk solution. Finally, it was incubated overnight at 4°C with an IRE1α antibody (Cell Signaling, #3294S) at 1:1000 dilution, followed by a two-hour incubation with horseradish peroxidase (HRP)-conjugated secondary antibodies at 1:10,000 dilution.

### Gel fractionation

100 μg of IRE1-KR-3P was crosslinked with DSS for 30 min on ice. The reaction was then loaded on Invitrogen 4-16% NativePAGE gel at 4°C for 4 h at 100V. Subsequently, the gel was cut in pieces at locations corresponding to monomer, dimer, and oligomer fractions. Gel fractions were then put in 100 μl of RNA cleavage buffer containing 4 μg of the T7 RNA transcript to digest overnight at 4°C. Finally, 10 μl of the reaction was used and run on a 3% agarose gel for visualization.

### Cell IRE1 Rescue

MDAMB231 or HCC1806 shIRE1α cell lines were transfected with a 2 kb IRE1α promoter-driven - shIRE1α resistant - GFP/His tagged IRE1α WT or R887A mutant - Neomycin resistant construct, using Mirus TransIT-X2 delivery system on 6 well plates. After 24h cells were transferred to individual T75 flasks for 4 days. Media was changed and cells were selected using Geneticin at 1.5mg/ml final for roughly 10 days, then FACS sorted for GFP positive cells.

### Viability Assay

4000 cells were plated in 4 replicates on 96-well Corning plates either standard flat clear-bottom or ULA (#7007). At plating, cells were treated with a 0.4 μg/ml Doxycycline (Clonetech) final concentration in 200 μl total volume. 7 days later, 100 μl of media was taken out very carefully from each plates, not disturbing the spheroids from the ULA plate. Cell viability was then assessed by CellTiter-Glo 3D, adding 100 μl of buffer (Promega #G9683) to each well and pipetting up and down a few times, and measured in a luminescence reader (Envision; PerkinElmer). The data depicted as Relative viability after Dox treatment is calculated from the means of quadruplicate samples, wherein viability of Dox treated cells is divided by that of untreated cells (ratio) and normalized to 2D mean viability.

## REFERENCES

1. Ron, D. & Walter, P. Signal integration in the endoplasmic reticulum unfolded protein response. Nat Rev Mol Cell Biol 8, 519–529 (2007).

2. Schroder, M. & Kaufman, R.J. The mammalian unfolded protein response. Annu Rev Biochem 74, 739–789 (2005).

3. Walter, P. & Ron, D. The unfolded protein response: from stress pathway to homeostatic regulation. Science 334, 1081–1086 (2011).

4. Hetz, C. The unfolded protein response: controlling cell fate decisions under ER stress and beyond. Nat Rev Mol Cell Biol 13, 89–102 (2012).

5. Shore, G.C., Papa, F.R. & Oakes, S.A. Signaling cell death from the endoplasmic reticulum stress response. Curr Opin Cell Biol 23, 143–149 (2011).

6. Tabas, I. & Ron, D. Integrating the mechanisms of apoptosis induced by endoplasmic reticulum stress. Nat Cell Biol 13, 184–190 (2011).

7. Bettigole, S.E. et al. The transcription factor XBP1 is selectively required for eosinophil differentiation. Nat Immunol 16, 829–837 (2015).

8. Chevet, E., Hetz, C. & Samali, A. Endoplasmic reticulum stress-activated cell reprogramming in oncogenesis. Cancer Discov 5, 586–597 (2015).

9. Grootjans, J., Kaser, A., Kaufman, R.J. & Blumberg, R.S. The unfolded protein response in immunity and inflammation. Nat Rev Immunol 16, 469–484 (2016).

10. Hetz, C., Chevet, E. & Harding, H.P. Targeting the unfolded protein response in disease. Nat Rev Drug Discov 12, 703–719 (2013).

11. Lin, J.H., Walter, P. & Yen, T.S. Endoplasmic reticulum stress in disease pathogenesis. Annu Rev Pathol 3, 399–425 (2008).

12. Chen, X. & Cubillos-Ruiz, J.R. Endoplasmic reticulum stress signals in the tumour and its microenvironment. Nature Reviews Cancer (2020).

13. Chen, X. et al. XBP1 promotes triple-negative breast cancer by controlling the HIF1alpha pathway. Nature 508, 103–107 (2014).

14. Harnoss, J.M. et al. IRE1α disruption in triple-negative breast cancer cooperates with anti-angiogenic therapy by reversing ER stress adaptation and remodeling the tumor microenvironment. Cancer Res (2020).

15. Harnoss, J.M. et al. Disruption of IRE1α through its kinase domain attenuates multiple myeloma. Proc Natl Acad Sci U S A 116, 16420–16429 (2019).

16. Urra, H., Dufey, E., Avril, T., Chevet, E. & Hetz, C. Endoplasmic Reticulum Stress and the Hallmarks of Cancer. Trends Cancer 2, 252–262 (2016).

17. Wang, M. & Kaufman, R.J. The impact of the endoplasmic reticulum protein-folding environment on cancer development. Nat Rev Cancer 14, 581–597 (2014).

18. Cox, J.S., Shamu, C.E. & Walter, P. Transcriptional induction of genes encoding endoplasmic reticulum resident proteins requires a transmembrane protein kinase. Cell 73, 1197–1206 (1993).

19. Lee, K.P. et al. Structure of the dual enzyme Ire1 reveals the basis for catalysis and regulation in nonconventional RNA splicing. Cell 132, 89–100 (2008).

20. Gardner, B.M. & Walter, P. Unfolded proteins are Ire1-activating ligands that directly induce the unfolded protein response. Science 333, 1891–1894 (2011).

21. Han, D. et al. IRE1alpha kinase activation modes control alternate endoribonuclease outputs to determine divergent cell fates. Cell 138, 562–575 (2009).

22. Korennykh, A.V. et al. The unfolded protein response signals through high-order assembly of Ire1. Nature 457, 687–693 (2009).

23. Shamu, C.E. & Walter, P. Oligomerization and phosphorylation of the Ire1p kinase during intracellular signaling from the endoplasmic reticulum to the nucleus. EMBO J 15, 3028–3039 (1996).

24. Tirasophon, W., Welihinda, A.A. & Kaufman, R.J. A stress response pathway from the endoplasmic reticulum to the nucleus requires a novel bifunctional protein kinase/endoribonuclease (Ire1p) in mammalian cells. Genes Dev 12, 1812–1824 (1998).

25. Hollien, J. et al. Regulated Ire1-dependent decay of messenger RNAs in mammalian cells. J Cell Biol 186, 323–331 (2009).

26. Hollien, J. & Weissman, J.S. Decay of endoplasmic reticulum-localized mRNAs during the unfolded protein response. Science 313, 104–107 (2006).

27. Maurel, M., Chevet, E., Tavernier, J. & Gerlo, S. Getting RIDD of RNA: IRE1 in cell fate regulation. Trends Biochem Sci 39, 245–254 (2014).

28. Brodsky, J.L. Cleaning up: ER-associated degradation to the rescue. Cell 151, 1163–1167 (2012).

29. Calfon, M. et al. IRE1 couples endoplasmic reticulum load to secretory capacity by processing the XBP-1 mRNA. Nature 415, 92–96 (2002).

30. Shen, X. et al. Complementary signaling pathways regulate the unfolded protein response and are required for C. elegans development. Cell 107, 893–903 (2001).

31. Yoshida, H., Matsui, T., Yamamoto, A., Okada, T. & Mori, K. XBP1 mRNA is induced by ATF6 and spliced by IRE1 in response to ER stress to produce a highly active transcription factor. Cell 107, 881–891 (2001).

32. Jurkin, J. et al. The mammalian tRNA ligase complex mediates splicing of XBP1 mRNA and controls antibody secretion in plasma cells. EMBO J 33, 2922–2936 (2014).

33. Lu, Y., Liang, F.X. & Wang, X. A synthetic biology approach identifies the mammalian UPR RNA ligase RtcB. Mol Cell 55, 758–770 (2014).

34. Peschek, J., Acosta-Alvear, D., Mendez, A.S. & Walter, P. A conformational RNA zipper promotes intron ejection during non-conventional XBP1 mRNA splicing. EMBO Rep 16, 1688–1698 (2015).

35. Hooks, K.B. & Griffiths-Jones, S. Conserved RNA structures in the non-canonical Hac1/Xbp1 intron. RNA Biol 8, 552–556 (2011).

36. Lhomond, S. et al. Dual IRE1 RNase functions dictate glioblastoma development. EMBO Mol Med 10 (2018).

37. Kimmig, P. et al. The unfolded protein response in fission yeast modulates stability of select mRNAs to maintain protein homeostasis. Elife 1, e00048 (2012).

38. Coelho, D.S. et al. Xbp1-independent Ire1 signaling is required for photoreceptor differentiation and rhabdomere morphogenesis in Drosophila. Cell Rep 5, 791–801 (2013).

39. Gaddam, D., Stevens, N. & Hollien, J. Comparison of mRNA localization and regulation during endoplasmic reticulum stress in Drosophila cells. Mol Biol Cell 24, 14–20 (2013).

40. Hollien, J. Evolution of the unfolded protein response. Biochim Biophys Acta 1833, 2458–2463 (2013).

41. Moore, K. & Hollien, J. Ire1-mediated decay in mammalian cells relies on mRNA sequence, structure, and translational status. Mol Biol Cell 26, 2873–2884 (2015).

42. Moore, K.A., Plant, J.J., Gaddam, D., Craft, J. & Hollien, J. Regulation of sumo mRNA during endoplasmic reticulum stress. PLoS One 8, e75723 (2013).

43. Oikawa, D., Tokuda, M., Hosoda, A. & Iwawaki, T. Identification of a consensus element recognized and cleaved by IRE1 alpha. Nucleic Acids Res 38, 6265–6273 (2010).

44. So, J.S. et al. Silencing of lipid metabolism genes through IRE1alpha-mediated mRNA decay lowers plasma lipids in mice. Cell Metab 16, 487–499 (2012).

45. Chang, T.K. et al. Coordination between Two Branches of the Unfolded Protein Response Determines Apoptotic Cell Fate. Mol Cell 71, 629–636 e625 (2018).

46. Lam, M., Marsters, S.A., Ashkenazi, A. & Walter, P. Misfolded proteins bind and activate death receptor 5 to trigger apoptosis during unresolved endoplasmic reticulum stress. Elife 9 (2020).

47. Lu, M. et al. Opposing unfolded-protein-response signals converge on death receptor 5 to control apoptosis. Science 345, 98–101 (2014).

48. Bae, D., Moore, K.A., Mella, J.M., Hayashi, S.Y. & Hollien, J. Degradation of Blos1 mRNA by IRE1 repositions lysosomes and protects cells from stress. J Cell Biol 218, 1118–1127 (2019).

49. Dufey, E. et al. Genotoxic stress triggers the activation of IRE1alpha-dependent RNA decay to modulate the DNA damage response. Nat Commun 11, 2401 (2020).

50. Tang, C.H. et al. Phosphorylation of IRE1 at S729 regulates RIDD in B cells and antibody production after immunization. J Cell Biol 217, 1739–1755 (2018).

51. Guydosh, N.R., Kimmig, P., Walter, P. & Green, R. Regulated Ire1-dependent mRNA decay requires no-go mRNA degradation to maintain endoplasmic reticulum homeostasis in S. pombe. Elife 6 (2017).

52. Prischi, F., Nowak, P.R., Carrara, M. & Ali, M.M. Phosphoregulation of Ire1 RNase splicing activity. Nat Commun 5, 3554 (2014).

53. Aragon, T. et al. Messenger RNA targeting to endoplasmic reticulum stress signalling sites. Nature 457, 736–740 (2009).

54. Belyy, V., Tran, N.H. & Walter, P. Quantitative microscopy reveals dynamics and fate of clustered IRE1alpha. Proc Natl Acad Sci U S A 117, 1533–1542 (2020).

55. Ghosh, R. et al. Allosteric inhibition of the IRE1alpha RNase preserves cell viability and function during endoplasmic reticulum stress. Cell 158, 534–548 (2014).

56. Li, H., Korennykh, A.V., Behrman, S.L. & Walter, P. Mammalian endoplasmic reticulum stress sensor IRE1 signals by dynamic clustering. Proc Natl Acad Sci U S A 107, 16113–16118 (2010).

57. Ricci, D. et al. Clustering of IRE1alpha depends on sensing ER stress but not on its RNase activity. FASEB J 33, 9811–9827 (2019).

58. Lopes, R., Agami, R. & Korkmaz, G. GRO-seq, A Tool for Identification of Transcripts Regulating Gene Expression. Methods Mol Biol 1543, 45–55 (2017).

59. Cross, B.C. et al. The molecular basis for selective inhibition of unconventional mRNA splicing by an IRE1-binding small molecule. Proc Natl Acad Sci U S A 109, E869–878 (2012).

60. Harrington, P.E. et al. Unfolded Protein Response in Cancer: IRE1alpha Inhibition by Selective Kinase Ligands Does Not Impair Tumor Cell Viability. ACS Med Chem Lett 6, 68–72 (2015).

61. Joshi, A. et al. Molecular mechanisms of human IRE1 activation through dimerization and ligand binding. Oncotarget 6, 13019–13035 (2015).

62. Tam, A.B., Koong, A.C. & Niwa, M. Ire1 has distinct catalytic mechanisms for XBP1/HAC1 splicing and RIDD. Cell Rep 9, 850–858 (2014).

63. Ferri, E. et al. Activation of the IRE1 RNase through remodeling of the kinase front pocket by ATP-competitive ligands. Nat Commun 11, 6387 (2020).

64. Korennykh, A.V. et al. Structural and functional basis for RNA cleavage by Ire1. BMC Biol 9, 47 (2011).

65. Niwa, M., Sidrauski, C., Kaufman, R.J. & Walter, P. A role for presenilin-1 in nuclear accumulation of Ire1 fragments and induction of the mammalian unfolded protein response. Cell 99, 691–702 (1999).

66. Shemorry, A. et al. Caspase-mediated cleavage of IRE1 controls apoptotic cell commitment during endoplasmic reticulum stress. Elife 8 (2019).

67. Zhang, Z. et al. TIPE1 induces apoptosis by negatively regulating Rac1 activation in hepatocellular carcinoma cells. Oncogene 34, 2566–2574 (2015).

68. Sundaram, A., Plumb, R., Appathurai, S. & Mariappan, M. The Sec61 translocon limits IRE1alpha signaling during the unfolded protein response. Elife 6 (2017).

69. Peschek, J. & Walter, P. tRNA ligase structure reveals kinetic competition between non-conventional mRNA splicing and mRNA decay. Elife 8 (2019).

70. Wu, T.D. & Watanabe, C.K. GMAP: a genomic mapping and alignment program for mRNA and EST sequences. Bioinformatics 21, 1859–1875 (2005).

71. Bendtsen, J.D., Nielsen, H., von Heijne, G. & Brunak, S. Improved prediction of signal peptides: SignalP 3.0. J Mol Biol 340, 783–795 (2004).

