## Supplementary material for "Noncanonical mRNA decay by the endoplasmic-reticulum stress sensor IRE1α promotes cancer-cell survival": gRIDD Supplemental Methods

### Supplemental method description of gRIDD algorithm

### Program and datasets

Our study empirically identified several examples of RIDD substrates, which we used to help refine the canonical pattern for RIDD cleavage sites. To this end, we used RNA transcripts that were cleaved by IRE-KR-0P as positive examples to be identified by our prediction rules, and transcripts cleaved only by IRE1-KR-3P as negative examples to be excluded.

To implement the prediction rules, we developed a computational program, called gRIDD (Genentech RIDD predictor), to scan a given transcript and report potential RIDD cleavage sites within it. The program first looks for candidate consensus sequences in the transcript. We used existing literature, which suggests that RIDD substrates have the consensus sequence CNGCNG, with four nucleotides highly conserved. The most common, or exact, consensus sequence has been observed to be CUGCAG. We allow for one mismatch from the exact consensus sequence, meaning that either position 2 or position 5 is permited to deviate, yielding the variant consensus sequences CxGCAG or CUGCxG, respectively, where x indicates a variation from the exact consensus sequence.

Canonical RIDD cleavage sites are believed to lie within a stem–loop structure, determined by the nucleotide content surrounding the consensus sequence. We therefore required the consensus sequence to reside within a loop of 6, 7, or 8 bp, where the nucleotides immediately outside the loop were complementary, meaning an A–U, G–C, or G–U pair. For a loop with 7 bp, we required the consensus sequence to be at the 5' end of the loop, leaving one additional nucleotide at the 3' section of the 6-nt consensus sequence. Likewise, for a loop with 8 bp, we required the consensus sequence to be at the 5' end of the loop.

For each candidate consensus sequence with a potential loop, our program then uses the RNAfold program to determine the secondary RNA structure surrounding the loop, providing that program with the constraint that the nucleotides immediately surrounding the loop be base-paired. To ensure that a stem–loop can exist in its global context, we provided RNAfold with a neighborhood of 55–60 nucleotides surrounding each candidate consensus sequence, testing each value from 60 down to 55, ending if and when a stem–loop structure surrounding the consensus sequence position was predicted. We used a range of nucleotides for the neighborhood length since we were uncertain about the boundaries where critical nucleotides at the end might have affected the global secondary structure.

The result of RNAfold is a secondary structure with minimum free energy, where bases are paired using the "(" and ")" symbols. Our program then analyzes the stem structure to require that it contain at least 3 consecutive base pairs. Each stem is further extended as far distally as possible, allowing for either a single mismatch at the same position in both legs of the stem, or a single bulge of up to 3 nucleotides. The total number of base pairs in the extended stem is used as a factor to discriminate RIDD sites from non-RIDD sites.

We applied the gRIDD program to the transcripts analyzed in our in-vitro experiments, which used a specified set of coding regions, from the start codon to the stop codon. The results of the program are shown in Table 1 below, with the positive examples used shown in black, and the negative ones in red. RNAs that failed to show cleavage by IRE1-KR-0P could potentially contain multiple sites, each of which constitutes a negative example of a RIDD site.

Table 1: Positive (black) and negative (red) examples from this study

| Gene | Transcript | Pos | Structure | Loop | Stem |
|---|---|---|---|---|---|
| **Observed 0P-type cleavage** | | | | | |
| *DGAT2* | BC015234.1 | 260 | ((((.(((((CUGCAGU))))).)))) | Exact | 9 |
| *XBP1#1* | NM_005080.3 | 519 | (((((((CUGCAGC))))))) | Exact | 7 |
| *BLOC1S1* | NM_001487.2 | 360 | ((((((CUGCAGU)))))) | Exact | 6 |
| *TGOLN2* | NM_006464.3 | 390 | ((((CUGCAGA)))) | Exact | 4 |
| *CD59* | NM_203331.2 | 78 | ((((CUGCAGU)))) | Exact | 4 |
| *PIGQ* | NM_004204.3 | 972 | ((.(((CUGCAG))).)) | Exact | 5 |
| *BMP4* | NM_130850.3 | 186 | ((((..(CUGCAG))))) | Exact | 5 |
| *XBP1#2* | NM_005080.3 | 484 | (((((CUGCuGA))))) | Var5 | 5 |
| **Observed 3P-type cleavage** | | | | | |
| *BCAM#1* | NM_005581.3 | 48 | ((((CUGCuG)))) | Var5 | 4 |
| *BCAM#2* | NM_005581.3 | 293 | ((((CcGCAGU)))) | Var2 | 4 |
| *MFAP2#1* | NM_002403.3 | 290 | ((((.(CUGCcG).)))) | Var5 | 5 |
| *MFAP2#2* | NM_002403.3 | 494 | (((CaGCAGCG))) | Var2 | 3 |
| *SIX2#1* | NM_016932.4 | 720 | (((CUGCcGU))) | Var5 | 3 |
| *SIX2#2* | NM_016932.4 | 66 | ((((((CaGCAGGG)))).)) | Var2 | 6 |
| *SNN* | NM_003498.5 | 108 | (((((.(CUGCgGCU).))))) | Var5 | 6 |
| *TNFAIP8L1* | NM_152362.1 | 219 | (((CUGCuGCG))) | Var5 | 3 |

We also applied the gRIDD program to positive examples of RIDD substrates from the literature, namely, a set of 13 human genes found to be RIDD substrates by Oikawa et al., 2010 (Table 2 below), and a set of 16 additional mammalian genes (4 human and 12 mouse) summarized from the literature by Maurel et al., 2014 (Table 3 below). For the Oikawa dataset, we used the full-length transcript from historical RefSeq archives that matched the transcript length given in their paper. For the Maurel genes, we analyzed all current transcript isoforms in RefSeq corresponding to the given gene. We considered examples from the literature as positive examples, although it is possible that other experiments may have confounded different cleavage modalities.

Table 2: Positive examples from Oikawa et al., 2010. Genes are colored in blue for subsequent reference. Ref indicates the cleavage site from the original paper.

| Gene | Transcript | Pos | Ref | Structure | Loop | Stem |
|---|---|---|---|---|---|---|
| *CCT3#1* | NM_001008883.1 | 1726 | | (((CUGCAGA))) | Exact | 3 |
| *CCT3#2* | NM_001008883.1 | 1754 | | ((((((((((CgGCAGU))).))))))) | Var2 | 10 |
| *FDFT1* | NM_004462.3 | 1073 | | ((((.(CaGCAG))))) | Var2 | 5 |
| *GEMIN5* | NM_015465.2 | 4862 | 4861 | (((((((((((CUGCAGA))))))))..))) | Exact | 11 |
| *IRAK1* | NM_001025242.1 | 2041 | | ((((((CUGCAGCU)))))) | Exact | 6 |
| *MKRN2#1*[a] | NM_014160.3 | 237 | | ((((((CUGCAGC)))))) | Exact | 6 |
| *MKRN2#2* | NM_014160.3 | 2249 | 2251 | ((((CaGCAGU)))) | Var2 | 4 |
| *MPC1*[b] | NM_016098.1 | 712 | | (((CUGCAGUA))) | Exact | 3 |
| *PDK2* | NM_002611.3 | 1978 | 1978 | (((((CUGCAGU))))) | Exact | 5 |
| *PEPD* | NM_000285.2 | 1455 | 1455 | ((((((((((.(CUGCAGC).)))))))))) | Exact | 11 |
| *PMF1* | NM_007221.2 | 756 | | (((((CaGCAGU).)))) | Var2 | 5 |
| *PPP2R1A* | NM_014225.3 | 1748 | 1748 | ((((CUGCAGA)))) | Exact | 4 |
| *PRKCD* | NM_006254.3 | 1736 | 1736 | ((((((((CUGCAGU).))))))) | Exact | 8 |
| *RUVBL1* | NM_003707.1 | 1536 | 1536 | ((((CUGCcGU)))) | Var5 | 4 |
| *YWHAQ* | NM_006826.2 | 994[c] | 1168[d] | ((((((.(CUGCAGC).)))))) | Exact | 7 |

**Induction of predictive rules**

We derived predictive rules for RIDD sites based on the results of our program, which reported the loop size and stem lengths for each candidate consensus sequence. We divided our analysis based on the three possible consensus patterns: exact (CUGCAG), variant 5 (CUGCxG), and variant 2 (CxGCAG). We found that the majority of known positive RIDD sites had an exact consensus sequence, followed by a smaller number with variant 5, and a smaller number with variant 2.

For the set of positive and negative examples with an exact consensus sequence, we categorized sites by their loop size and stem lengths (Table 4 below). Genes are colored according to their source, using the same colors as in Tables 1–3. Human genes are shown in all capital letters, while mouse genes are shown with the first letter capitalized, according to standard convention. The most prevalent loop size was 7 nt, with some observed to have loop sizes of 6 or 8 nt. In all cases, the positive examples had stem lengths of 3 or more bp. Therefore, we can capture known RIDD sites around exact consensus sequences by allowing for loop sizes of 6, 7, or 8 bp and requiring 3 or more base pairs in their extended stems.

Table 3: Positive examples from Maurel et al., 2014. Genes are colored in light blue for subsequent reference. Genes from Oikawa et al., 2010, are excluded, as are genes that have no entry as an NM or XM transcript in RefSeq: 28S RNA, MIR17, and $\mu$s.

| Gene[a] | Transcript | Pos | Structure | Loop | Stem |
|---|---|---|---|---|---|
| *Angptl3* | NM_013913.4 | 204 | ((((.(CUGCAGCU))))) | Exact | 5 |
| *Bloc1s1* | NM_015740.3 | 449 | (((((CUGCAGU)))).)) | Exact | 6 |
| *Col6a1* | NM_009933.4 | 1945 | ((((((CUGCuGU))))))) | Var5 | 7 |
| *Cyp2e1* | NM_021282.3 | 1326 | ((((((CUGCAGG))))))) | Exact | 7 |
| *ERN1* | XM_017024347.2 | 3086 | (((((CUGCAGG)))))) | Exact | 6 |
| *GPC3* | NM_004484.4 | 1374 | (((((..(CUGCAGCC)))))) | Exact | 6 |
| *Galnt2* | NM_139272.2 | 589 | ((((((CUGCgGA))).)))) | Var5 | 7 |
| *Hgsnat#1* | NM_029884.1 | 159 | (((((..((CUGCuGC))))))) | Var5 | 7 |
| *Hgsnat#2* | NM_029884.1 | 752 | (((((((.(CaGCAGA).))))))) | Var2 | 8 |
| *Itgb2* | NM_008404.5 | 1250 | (((((CUGCAGU)).)))) | Exact | 6 |
| *PER1* | NM_002616.3 | 18 | (((((CUGCgGG)).))) | Var5 | 5 |
| *Pdgfrb* | NM_008809.2 | 4940 | (((CaGCAGC))) | Var2 | 3 |
| *Pmp22* | NM_001302257.1 | 169 | (((CUGCAGGC))) | Exact | 3 |
| *RTN4* | NM_020532.5 | 2663 | (((CUGCAGUU))) | Exact | 3 |
| *Scara3* | NM_172604.3 | 219 | (((.(((((CUGCAGA))))).))) | Exact | 8 |
| *Tapbp* | NM_009318.2 | 591 | (.((((CUGCuGG))))) | Var5 | 5 |

[a] No stem–loop structures with an endomotif were found for Ces1 (Ces1a, Ces1b, or Ces1c), Cyp1a2, or PDIA4.

Table 4: Exact consensus sequences: CUGCAG

| Loop size | Stem ≥ 3 | Stem < 3 |
|---|---|---|
| 6 | *BMP4*(5), *PIGQ*(5) | |
| 7 | *GEMIN5*(11), *PEPD*(11), *DGAT2*(9), *PRKCD*(8), *Scara3*(8), *XBP1#1*(7), *YWHAQ*(7), *Cyp2e1*(7), *BLOC1S1*(6), *Bloc1s1*(6), *ERN1*(6), *Itgb2*(6), *MKRN2#1*(6), *PDK2*(5), *CD59*(4), *TGOLN2*(4), *PPP2R1A*(4) | |
| 8 | *GPC3*(6), *IRAK1*(6), *Angptl3*(5), *MPC1*(3), *Pmp22*(3), *RTN4*(3) | |

Sites with variant 5 consensus sequences are categorized similarly in Table 5 below. Here, we have a negative example from our study (*SIX2*, site #1) that suggests that for a loop size of 7 nt, 4 or more bp are required in the stem. Other negative examples from our study suggest that, even with stems of 4 or more bp, loop sizes of 6 or 8 bp are not substrates for 0P-type cleavage. Therefore, it appears that the predictive rules for variant 5 consensus sequences allow for only a loop size of 7 nt and requires 4 or more base pairs in their extended stems.

Table 5: Variant 5 consensus sequences: CUGCxG

| Loop size | Stem ≥ 4 | Stem < 4 |
|---|---|---|
| 6 | *MFAP2#1*(5), *BCAM#1*(4) | |
| 7 | *Col6a1*(7), *Galnt2*(7), *Hgsnat#1*(7), *XBP1#2*(5), *PER1*(5), *Tapbp*(5), *RUVBL1*(4) | *SIX2#1*(3) |
| 8 | *SNN*(6) | *TNFAIP8L1*(3) |

Finally, for sites with variant 2 consensus sequences, Table 6 below shows positive examples from our study and the literature for a loop size of 7 nt with stem lengths of 5 or more bp. *BCAM* site #2 suggests that a stem length of 4 bp is not sufficient when a variant 2 consensus sequence is present, and *SIX2* site #2 also suggests that a loop size of 8 nt is not a substrate for IRE1-KR-0P type cleavage with a variant 2 consensus sequence. The example *FDFT1* from the literature can be accounted for by allowing a loop size of 6 nt with a variant 2 consensus sequence. However, we do not wish to implement a rule for a single example from the literature. Therefore, a predictive rule for variant 2 consensus sequences appears to allow only for a loop size of 7 nt and requires 5 or more base pairs in their extended stem.

Table 6: Variant 2 consensus sequences: CxGCAG

| Loop size | Stem ≥ 5 | Stem < 5 |
|---|---|---|
| 6 | *FDFT1*(5) | |
| 7 | *CCT3*(10), *Hgsnat#2*(8), *PMF1*(5) | *BCAM#2*(4), *MKRN2#2*(4), *Pdfgrb*(3) |
| 8 | *SIX2#2*(6) | *MFAP2#2*(3) |

**Results**

Our predictive rules are able to predict all positive examples and to exclude all negative examples from Table 1. In addition, our program can predict 12 of the 13 genes reported by Oikawa et al., 2010, missing *FDFT1* and showing a different cleavage site for *MKRN2*. Our program identifies 12 of the 16 additional mammalian sites reported by Maurel et al., 2014, missing *Pdgfrb*, as well as *Ces1*, *Cyp1a2*, and *PDIA4*, for which no valid consensus sequences with a stem–loop structure was found in the available RefSeq transcripts. To test our rules further, we selected four additional genes with evidence of cleavage in the presence of *IRE1*: *AIM2*, *CCDC69*, *GBA*, and *WT1*. The predictions of our program and experimental results are shown in Table 7 below. In all four cases, experimental results were consistent with our predictive rules.

Table 7: Test cases

| Gene | Transcript | Pos | Structure | Loop | Stem | Pred | Obs |
|---|---|---|---|---|---|---|---|
| *AIM2* | NA | | | | | | 3P |
| *CCDC69* | NM_015621.2 | 786 | (((CaGCAG))) | Var2 | 3 | | 3P |
| *GBA* | NM_000157.4 | 612 | (.(((CUGCAGUU))).) | Exact | 4 | 0P | 0P |
| *WT1* | BC032861.2 | 120 | (((((((((CaGCAGG))))).)))) | Var2 | 9 | 0P | 0P |

Our rules could be made more sophisticated by using quantitative criteria, such as the minimum free energy predicted for the stem–loop structures. However, it is not clear whether minimum free energy is the most appropriate criterion; alternative stem–loop conformations could suffice, even if not at the minimum but still highly probable. Nevertheless, the number of base pairs in the stem serves as a proxy for energy calculations, with more base pairs helping to stabilize the stem–loop structure. Other considerations of secondary structure may also play a role, such as the presence of an additional hairpin loop upstream of the RIDD cleavage site. Such hairpins have been found to affect ribosomal processing, resulting in translational stalling. However, it is not clear if such hairpins would also affect in vitro cleavage.

Our analysis suggests that a tradeoff exists between the consensus sequence and the stem–loop structure, with an exact consensus sequence allowing for variable loop sizes and shorter stems, while the variant 2 and variant 5 consensus sequences require a loop size of 7 nt and greater stem lengths. This tradeoff suggests that RIDD cleavage depends on both the consensus sequence and the stem–loop structure as interrelated factors.
